# Geogenomic predictors of genetree heterogeneity in an Amazonian bird (*Thamnophilus aethiops*)

**DOI:** 10.1101/2023.08.22.554279

**Authors:** Lukas J. Musher, Glaucia Del-Rio, Rafael S. Marcondes, Robb T. Brumfield, Gustavo A. Bravo, Gregory Thom

**Affiliations:** The Academy of Natural Sciences of Drexel University, Department of Ornithology, Philadelphia, PA, USA; Cornell Laboratory of Ornithology and Department of Ecology and Evolutionary Biology, Cornell University, Ithaca, NY, USA; Department of Biology and Museum of Natural Science, Louisiana State University; Department of BioSciences, Rice University, USA; Museum of Natural Science and Department of Biological Sciences, Louisiana State University, Baton Rouge, LA 70803, USA; Sección de Ornitología, Colecciones Biológicas, Instituto de Investigación de Recursos Biológicos Alexander von Humboldt, Claustro de San Agustín, Villa de Leyva, Boyacá, Colombia; Museum of Comparative Zoology and Department of Organismic and Evolutionary Biology, Harvard University, Cambridge, MA, USA

**Keywords:** Amazon, biogeography, gene flow, genome architecture, geogenomics, neural network, reproductive isolation, speciation

## Abstract

Can knowledge about genome architecture inform biogeographic and phylogenetic inference? Selection, drift, recombination, and gene flow interact to produce a genomic landscape of divergence wherein patterns of differentiation and genealogy vary nonrandomly across the genomes of diverging populations. For instance, genealogical patterns that arise due to gene flow should be more likely to occur on smaller chromosomes, which experience high recombination, whereas those tracking histories of geographic isolation (reduced gene flow caused by a barrier) and divergence should be more likely to occur on larger and sex chromosomes. In Amazonia, populations of many bird species diverge and introgress across rivers, resulting in reticulated genomic signals. Herein, we used reduced representation genomic data to disentangle the evolutionary history of four populations of an Amazonian antbird, *Thamnophilus aethiops,* whose biogeographic history was associated with the dynamic evolution of the Madeira River Basin. Specifically, we evaluate whether a large river capture event ca. 200 kya, gave rise to reticulated genealogies in the genome by making spatially explicit predictions about isolation and gene flow based on knowledge about genomic processes. We first estimated chromosome-level phylogenies and recovered two primary topologies across the genome. The first topology (T1) was most consistent with predictions about population divergence, and was recovered for the Z chromosome. The second (T2), was consistent with predictions about gene flow upon secondary contact. To evaluate support for these topologies, we trained a convolutional neural network to classify our data into alternative diversification models and estimate demographic parameters. The best-fit model was concordant with T1 and included gene flow between non-sister taxa. Finally, we modeled levels of divergence and introgression as functions of chromosome length, and found that smaller chromosomes experienced higher gene flow. Given that (1) gene-trees supporting T2 were more likely to occur on smaller chromosomes and (2) we found lower levels of introgression on larger chromosomes (and especially the Z-chromosome), we argue that T1 represents the history of population divergence across rivers and T2 the history of secondary contact due to barrier loss. Our results suggest that a significant portion of genomic heterogeneity arises due to extrinsic biogeographic processes such as river capture interacting with intrinsic processes associated with genome architecture. Future biogeographic studies would benefit from accounting for genomic processes, as different parts of the genome reveal contrasting, albeit complementary histories, all of which are relevant for disentangling the intricate geogenomic mechanisms of biotic diversification.

## Introduction

A key goal of speciation research is to understand the biogeographic mechanisms associated with population divergence and homogenization (Endler 1977). Although the reduction of gene flow between populations caused by biogeographic barriers (henceforth geographic isolation) typically plays a crucial role in speciation, populations may diverge despite high gene flow (Nosil 2008). Such gene flow, especially between non-sister populations, can result in heterogeneous levels of differentiation across the genome (Keller et al. 2013; Gompert et al. 2014; Mallet et al. 2016; Meier et al. 2017; Pulido-Santacruz et al. 2020), a pattern further intensified by interactions with selection and genome architecture (Feder et al. 2012; Cruickshank and Hahn 2014; Irwin et al. 2018; Manthey et al. 2021). For example, genomic regions experiencing strong disruptive selection or low recombination (e.g., large chromosomes (Haenel et al. 2018) may resist the homogenizing effects of gene flow, resulting in elevated peaks of divergence and contrasting genealogies when compared with other parts of the genome. Thus, the genomic landscape of divergence not only reflects disparate levels of differentiation between populations but also a profusion of genealogical relationships (Fontaine et al. 2015; Mallet et al. 2016; Wen et al. 2016). Dealing with and modeling evolutionary history in light of this heterogeneous landscape is crucial for obtaining a detailed understanding of biogeographic processes (Provost et al. 2022).

Although the genomic landscape can be highly heterogeneous, the signals of distinct processes, such as divergence and gene flow, are nonrandomly distributed across the genome (Van Doren et al. 2017). For instance, variation in recombination rates directly impacts levels of gene flow and divergence (Wang et al. 2022). One widely recognized mechanism by which this operates is the breakdown of blocks of linked loci affected by selection. Specifically, linked selection (selection in the genome impacting nearby sites) can significantly diminish genetic variation in blocks of linked loci through genetic hitchhiking of nearby sites (Lohmueller et al. 2011; Feder et al. 2012). When linked selection within populations is strong, measures of relative divergence, such as F_ST_, which contain a term for within population variation, are expected to increase, but absolute divergence, d_xy_, may be reduced or unaffected by the corresponding depletion of allelic diversity (Charlesworth 1998; Nachman and Payseur 2012; Cruickshank and Hahn 2014; Van Doren et al. 2017; Irwin et al. 2018). In regions of the genome where recombination is high, blocks of linked loci are more frequently broken down, lessening the effects of linked selection (Tigano et al. 2022). Where recombination is low, however, the effects of selection are elevated as longer blocks of linked loci are able to persist. Recombination rate varies considerably across the genome and is particularly associated with chromosome size. Because each chromosome must undergo at least one crossing-over event during Meiosis (Mather 1938), smaller chromosomes experience more recombination per base than large chromosomes (Haenel et al. 2018; Tigano et al. 2022). Thus, regions of the genome with higher rates of recombination, such as smaller chromosomes, are also expected to have higher rates of introgression, when population divergence occurs with gene flow (Manthey et al., 2021; Martin, Davey, Salazar, & Jiggins, 2019). Quantifying the variation and predictability of these genomic processes that are intrinsic to organisms can help shed light on the reticulated history of recent radiations.

In contrast to intrinsic genomic architecture, biogeographic history is an important extrinsic factor influencing reticulation and the genomic landscape (Burbrink and Gehara 2018; Thom et al. 2021; Provost et al. 2022). As levels of isolation associated with physiographic barriers vary through space and time, so too do rates of selection, gene flow, and divergence (Endler 1977; Aguilée et al. 2013; Delmore et al. 2018; He et al. 2019). In many parts of the world, population isolation and connectivity vary, in part, as functions of spatiotemporal variation in the environment (Flantua et al. 2019; He et al. 2019; Musher et al. 2019). For example, rates of isolation and gene flow among populations that diverged across the Isthmus of Panama may have been affected by the wax and wane of humid and dry forests across that region (David Webb 1991; Vrba 1992; Smith et al. 2012; Musher et al. 2020). Likewise, sea-level fluctuations and rainfall patterns have directly affected the distribution and amount of flooded forest habitat in Amazonia, which also affected levels of gene flow between populations of organisms that occur there (Thom et al. 2020; Sawakuchi et al. 2022; Luna et al. 2023). In Amazonian lowlands, differentiated populations often experience pervasive gene flow, sometimes from multiple non-sister lineages, a factor that complicates inference about their historical relationships, biogeography, and systematics (Pulido-Santacruz et al. 2018; Del-Rio et al. 2022; Musher et al. 2022). This is because gene flow between non-sister taxa results in a network of interpopulation relationships that violates the assumptions of a bifurcating model of evolutionary history (Mallet et al. 2016; Thom et al. 2018). Thus, if biogeography drives opportunities for isolation and contact between non-sister taxa, it should result in distinct predictable signatures of genealogy in the genomes of diverging populations.

Many lowland *terra-firme* (non-flooded forest) Amazonian birds have geographically isolated populations across rivers, yet experience high levels of gene flow (Barrera-Guzmán et al. 2022; Del-Rio et al. 2022; Musher et al. 2022). Rivers are key biogeographic barriers for many lowland Amazonian birds, driving population isolation and genetic structure across the landscape (Sick 1967; Capparella 1991; Ribas et al. 2012; Smith et al. 2014; Ferreira et al. 2017). However, three well-known features of Amazonian lowlands add complexity to this system. First, the Amazon Basin, especially its southern portion, is characterized by several large tributaries running in quasi-parallel, forming isolated blocks of habitat (interfluves) wherein a given taxon may be surrounded by two or more closely related taxa that occur on opposite river margins. Second, Amazonian rivers get narrower toward their headwaters, which is associated with increased gene flow across their upper portions (Weir et al. 2015). Finally, lowland river basins continuously rearrange via tributary capture (the movement of a tributary from one basin to another) and avulsion (the erosion of channel boundaries, leading to channel migration) (Gascon et al. 2000; Albert et al. 2018). In this geographic configuration, there are opportunities for multiple non-sister taxa to interact, and partially isolated populations can experience gene flow across rivers, leading to highly reticulated patterns of diversification across species’ genomes (Musher et al. 2022).

Inferring the history of population isolation and gene flow under these conditions is a major challenge for researchers studying Amazonia because limited knowledge about the historical relationships of taxa also hampers an understanding of the mechanisms that contribute to the region’s high biodiversity (Cracraft et al. 2020). Previous studies of Amazonian birds have greatly advanced our knowledge of the history of the region’s taxa, but in many cases, limited data or sparse spatial sampling has resulted in weak resolution of species’ complex histories of isolation and gene flow. Genomic approaches, however, are revealing many of the evolutionary and biogeographic mechanisms driving species accumulation in the Neotropics (Thom et al. 2018, 2020; Pulido-Santacruz et al. 2020; Schley et al. 2020). This is especially important given geologists’ growing understanding that the Amazonian landscape has been highly dynamic (Bicudo et al. 2019; Pupim et al. 2019; Ruokolainen et al. 2019). Thus, important questions for Amazonian biogeography include, (1) how do we infer population history under conditions of high gene flow among non-sister taxa, especially in the context of a dynamic landscape? (2) What are the consequences of these complex histories of isolation and gene flow for Amazonian organisms at the genomic level? (3) do genealogical patterns vary predictably across the genome in a way that is informative for biogeographic inference? And (4) to what extent can reduced representation genomic data shed light on these questions?

In this study, we address these questions by testing competing hypotheses for the biogeographic history of a passerine bird, the White-shouldered antshrike *Thamnophilus aethiops* (Thamnophilidae), in southern Amazonia. The southern Amazonian lowlands are particularly dynamic, and experienced a major riverine restructuring, wherein large tributaries moved among watersheds and across the landscape during the Quaternary (Latrubesse 2002; Rossetti 2014; Ruokolainen et al. 2019; Rossetti et al. 2021). Some of these past movements occurred near the headwaters of the modern Madeira River, a large tributary of the Amazon that is a well-known biogeographic barrier for many bird species (Fernandes 2013; Smith et al. 2014; Silva et al. 2019). For example, paleochannels between the modern upper Madeira and Purus Rivers indicate that the Madeira extended its basin by capturing a large tributary of the Purus ca. 200 kya or less (Fig. 1) (Ruokolainen et al. 2019). This suggests that the upper portion of the Madeira formed more recently than the lower, likely becoming a barrier for many terrestrial organisms over two stages.

**Figure 1:**
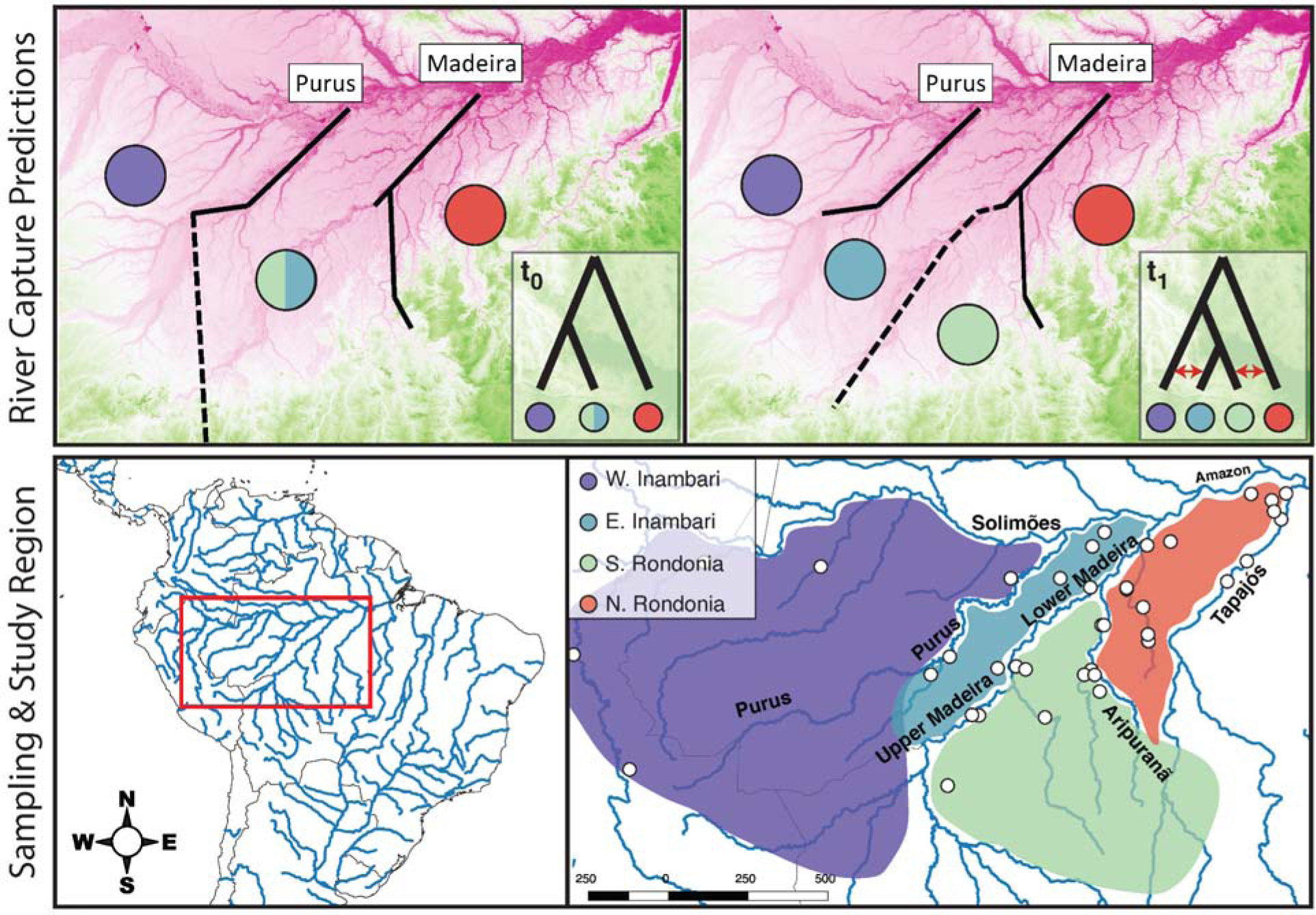
Spatially explicit predictions of the river capture scenario (top row). The stippled lines show the movement of a paleo-river, historically draining water via the Purus watershed (time=t_0_) and its subsequent capture by the modern Madeira watershed (time=t_1_). Phylogenetic predictions assume a set of historical relationships (t_0_); the predicted topology (t_1_) matches that inferred from mitochondrial DNA in a previous study (Thom and Aleixo, 2015). Red arrows indicate opportunities for gene flow between non-sister populations. The maps in the top row are colored by topography, with dark pinks representing lower elevations (e.g. river channels) and darker greens representing uplands. The bottom row shows the study region, sampling (white circles), and key interfluvial areas: W Inambari (purple), E Inambari (teal), S Rondônia (pale green), and N Rondônia (red).

The Madeira River capture scenario provides predictions about the history of population divergence and gene flow, wherein populations east of the Upper Madeira (Southern Rondônia; Fig. 1) are expected to be more closely related to populations west of the Madeira (Inambari) than to other populations within Rondônia (Fernandes 2013; Ferreira et al. 2017). Such a scenario also suggests that there will be more genetic differentiation, and therefore potential incompatibilities across the genomes of more deeply diverged populations (e.g., those in Rondônia) than between more recently diverged populations (e.g., populations within Inambari). Previous studies of *T. aethiops* have been equivocal with respect to these predictions (Thom and Aleixo 2015; Musher et al. 2022). For example, the mitochondrial (mtDNA) phylogeny is consistent with the expectations of river capture, recovering a sister relationship between populations on opposite sides of the Madeira River (but not its downstream portions). Genomic data were more ambiguous, however, recovering relatively strong support for the monophyly of Rondônia and Inambari populations, which conflicts with the expectations of river capture. Thus, if the population in S Rondônia is historically related to Inambari populations, as the river capture scenario predicts, then we expect regions of the genome supporting monophyly of Rondônia to be driven by introgressive hybridization after secondary contact. Herein, we test these alternative genealogical predictions while also utilizing and modeling information about genome architecture. For instance, if gene flow increases on smaller autosomes, genealogies resulting from secondary contact should be less likely on larger autosomes and sex chromosomes. Rather, regions of the genome inferred to have low gene flow should better track phylogeny *sensu stricto* (i.e., the history of population divergences).

## Methods

### Study taxon

*Thamnophilus aethiops* is well-suited for testing our hypotheses as it exhibits both subspecific variation and genetic structure across southern Amazonian rivers (Thom and Aleixo 2015; Musher et al. 2022). Populations east of the Madeira River belong to a single subspecies, *T. a. punctuliger*. Populations west of the Madeira fall into one of three subspecies: *T. a. injunctus* occurs between the Purus and Madeira Rivers, *T. a. juruanus* occurs between the Jurua and Purus Rivers, and *T. a. kapouni* occurs west of the Juruá. Superficially, the phenotypes of *T. a. injunctus* and *T. a. punctuliger* are most similar, with individuals of both taxa being lighter gray overall and marked with white spots on the wing coverts, unlike populations farther west. Moreover, *T. aehiops* is a fairly sedentary understory species, so gene flow expectations across stable rivers are relatively low.

### Sampling and GBS data assembly

We downloaded genotyping-by-sequencing (GBS) data from a previous study (Musher et al. 2022), and re-assembled it using iPyrad version 0.9.81 (Elshire et al. 2011; Eaton and Overcast 2020). These data are available on the NIH Sequence Read Archive under project ID PRJNA966941. We were interested in the history and relationships of populations west of the Tapajós River and south of the Amazon River (southwestern Amazon Basin), so we excluded samples falling outside of this region. This dataset totaled 51 samples (Tables S1 & S2). First, we demultiplexed and cleaned raw reads. Specifically, we used TGCA as our restriction overhang (Pstl-I, which cleaves DNA at 5′-CTGCA/G-3′ sites was the restriction enzyme used in library prep), allowed a maximum of five low-quality (Q<20) base calls in each read with a Phred Q-score offset of 33, and discarded any cleaned reads with fewer than 35 bp. We then mapped cleaned reads to a reference genome of a closely-related species *Thamnophilus caerulescens* (divergence time: ∼4 Ma; Harvey et al. 2020; see Supplemental Materials). iPyrad uses vsearch (Rognes et al. 2016) to merge overlapping paired reads and then bwa (Li and Durbin 2009) to map those paired reads to a reference genome and determine locus homology, discarding any unmapped or replicated reads. In sum, paired-end reads are mapped to the reference genome based on sequence similarity and reads that cluster at the same genomic loci are then aligned using muscle (Edgar 2022). Each of these “aligned clusters” represents a single GBS locus. We applied a minimum statistical coverage of six, which is considered the minimum value for accurate base calling that is widely used in RADSeq studies (Eaton and Overcast 2020; Barreto et al. 2022; Donoghue et al. 2022; Hanes et al. 2022). When estimating consensus allele sequences from clustered reads, we allowed default settings of a maximum of 5% ambiguous base calls and 5% heterozygous sites. These last two filters reduce the risk of incorporating erroneous alignments into the dataset which are likely to have increased heterozygosity and may be caused when many ambiguous bases are included in a consensus.

### Population structure and ancestry

Characterization of the population structure of *T. aethiops* is crucial for delineating populations for downstream analysis and characterization of patterns of admixture. We first investigated genetic structure among populations using principal components analysis (PCA), implemented in the iPyrad API (Eaton and Overcast 2020). Specifically, we used *k-*means iterative clustering to assign individuals to populations under a four-population model (*k*=4). We iteratively clustered sample single-nucleotide polymorphisms (SNPs) present across 90% of individuals and clustered individuals based on an assumed value for the number of populations. We repeated this clustering five times, allowing more missing data at each successive iteration until reaching a minimum coverage of 75% of individuals for a given SNP. Next, missing genotypes for all samples were imputed by choosing one random genotype from the assigned population based on the *k-*means assignments. This method allowed imputation without *a priori* geographic bias. We then performed PCA on unlinked genotype calls (one randomly sampled genotype per locus) in this imputed dataset.

We then used sparse non-negative matrix factorization (sNMF), implemented in the R package LEA3 (Frichot et al. 2014; Gain and François 2021), to calculate admixture coefficients for each population and infer the best-fit number of ancestral populations (*k*) in the dataset. We tested values of *k* from one through five to infer the optimal number of ancestries. As sNMF results are sometimes sensitive to the regularization parameter, α, we explored our results under multiple values of α (10,100,1000,10000) using only ten replicates. However, given our relatively large dataset, our results were stable across all α-values, so we performed 1000 iterations of each *k* value using α=10 to obtain our final results. The best-fit value of *k* was determined by choosing the *k* with the lowest cross-entropy value (Frichot et al. 2014). In addition to the best-fit value of *k*, we also visualized the results from other values of *k*, to better characterize additional, biologically relevant population structure present in the data. To assess the stability of population assignments in sNMF, we ran this procedure multiple times, replicating the 1000 iterations of sNMF over multiple runs.

### Genetrees, chromosome trees, and species tree topologies

We evaluated the phylogenetic relationships for the populations recovered in sNMF and PCA analyses (K=4), which were also consistent with previously published mitochondrial clades (Thom and Aleixo 2015). We estimated genetrees for sliding windows across the genome using TreeSlider, a python program available within the iPyrad API (Eaton and Overcast 2020). TreeSlider keeps the most common allele for each locus in each population and estimates the maximum likelihood phylogeny at each locus using RAxML (Stamatakis 2014). We specifically compared the results from TreeSlider for 10kb, 50kb, 100kb, and 200kb sliding windows, first requiring a minimum of five SNPs, and then a minimum of ten SNPs to retain each window. We found that 50kb windows with a minimum of five SNPs retained the most (4,858; Table S2) loci, so we used the genetrees from this dataset for all downstream phylogenetic analyses.

To understand how phylogenetic history varied across the genome and among chromosomes, we analyzed genetree results at multiple scales using ASTRAL v5.7.8 (Zhang et al. 2018). ASTRAL estimates an unrooted species tree given a set of genetrees by identifying the maximum number of shared induced quartets within the provided genetrees. We used ASTRAL to estimate the interrelationships of four populations of interest, corresponding to those identified using sNMF and PCA. To understand how these relationships varied among parts of the genome, we first ran ASTRAL for all 4,858 genetrees together (genome-wide species tree), second for only autosomal genetrees (autosomal tree; i.e., excluding genetrees from sex chromosomes), and finally for genetrees from each chromosome independently (chromosomal trees). As no loci on Chromosome 33 met our criteria (50kb minimum of 5 SNPs), we excluded this chromosome from all ASTRAL analyses.

To quantify variation in support for competing phylogenetic hypotheses, we calculated unrooted topology weights for alternative topologies across windows of the genome using TWISST (Martin and Van Belleghem 2017). This approach provides an assessment of the relative likelihood of alternative topologies for individual genomic windows. Here, we estimated genetrees from SNPs on sliding windows using PHYML v3.0 (Guindon et al. 2010) following Martin and Van Belleghem (2017). Due to the patchiness of GBS datasets, we analyzed windows of varying length with exactly 100 SNPs each. The larger window size and stricter filtering applied here enabled the estimation of relatively high-resolution genetrees, reducing noise in this analysis. We performed two independent runs testing the relationships between 4 taxa. First, we calculated topology weights for three unrooted topologies using the 4 ingroup populations. This configuration allowed us to test the sister relationship between E and W Inambari populations. Next, we added the reference genome (*T. caerulescens*) to the dataset and removed the W Inambari population, allowing us to test S Rondônia as sister to Inambari populations. Because TWISST quantifies the relative weights of unrooted topologies, running our analyses with four taxa enabled us to evaluate support for alternative quartets identified in ASTRAL as well as the mitochondrial topology, without the noise of more complicated five-taxon statements. Initial tests with our data set revealed that 5 taxon runs had poor resolution given the larger set of alternative topologies.

Given the overall low support for the mtDNA tree across the genome (see Results) we explored whether the few genomic windows with high support for mtDNA topology were linked to nuclear genes associated with mitochondrial activity. We selected genes associated with mitochondrial activity according to Morales et al. (2018) (734 N-mt genes; Gene Ontology term: 0005739 and 0006119) using our annotated reference genome, located within 100 kb of windows with topology weights above three standard deviations from the mean for the mtDNA tree.

### Demographic modeling

To estimate genome-wide topology for the four populations while accounting for gene flow, we used a combination of coalescent simulations performed with PipeMaster (Gehara et al. 2017), and supervised machine learning implemented in Keras v2.3 (Arnold 2017; https://github.com/rstudio/keras). We first simulated data under three demographic models matching alternative phylogenetic hypotheses under the effect of gene flow between selected populations. For the purposes of this section, we abbreviate N Rondônia as “NR,” S Rondônia as “SR,” W Inambari as “WI,” and E Inambari as “EI.” The simulated models were: Model 1) (NR,(SR(EI, WI))) with gene flow between SR and both NR and EI (M_SR<->NR_ and M_SR<->EI_); Model 2) ((NR, SR),(EI, WI)) with gene flow between SR and EI (M_SR<->EI_); and Model 3) (NR,(WI,(EI, SR))) with gene flow between populations on both sides of the Madeira River (M_SR<->NR_ and M_WI<->EI_). Each of the three models matches the results obtained with independent phylogenetic analysis: Models 1 and 2 match the alternative topologies obtained with ASTRAL (T1 and T2; see Results); whereas Model 3 matches the mtDNA topology of Thom and Aleixo (2015) (T3; see Results). Symmetric gene flow was allowed in the model between pairs of non-sister populations potentially in contact (e.g., occurring on opposite sides of a river) that could explain the discordance observed in phylogenetic estimates. We only allowed symmetric gene flow between populations to reduce the number of parameters of our models and because our primary goal was to test for alternative phylogenetic topologies while accounting for gene flow; our intent was not to estimate the absolute values of demographic parameters. For all models we set relatively large and uniform priors based on reasonable values for lowland Amazonian species: Ne 100,000 – 1,000,000 diploid individuals (for all populations); M_pop1<->pop2_ 0.1 - 3.0 migrants per generation; Tdiv (Divergence time in generations) Model 1: WI/EI 50,000 - 800,000, SR/WI+EI 500,000 - 1,200,000, NR/WI+EI+SR 750,000 - 1,500,000; Model 2: WI/EI 50,000 - 800,000, NR/SR 50,000 - 800,000, NR+SR/WI+EI 750,000 - 1,500,000; and Model 3: EI/SR 50,000 - 800,000, WI/EI+SR 500,000 - 1,200,000, NR/WI+EI+SR 750,000 - 1,500,000. We assumed a fixed mutation rate of 2.42 × 10^−9^ mutations per generation and a one-year generation time (Jarvis et al. 2014; Zhang et al. 2014). Although an assumed one-year generation time may underestimate the true generation time, limited data about survivorship and reproduction in Amazonian birds precludes a more precise estimate (Saether et al. 2005).

To obtain the observed data on which simulations were based, we initially converted the alleles file produced by iPyrad into individual fasta alignments with the iPyrad.alleles.loci2fasta function of PipeMaster. Since PipeMaster is sensitive to missing data, we also applied multiple filters to the sequence data. First, we removed individuals missing more than 50% of the loci from alignments using Alignment_Refiner_v2.py (Portik et al. 2016) and excluded alignment positions not recovered for at least 80% of the individuals using trimAL (Capella-Gutierrez et al. 2009). Lastly, we excluded loci with fewer than 51 individuals, shorter than 100bp, and missing more than 50% of the sites using AMAS and custom R scripts (Borowiec 2016). The genetic data for each model was simulated on PipeMaster based on the number of retained loci, matching their length and number of individuals using msABC (Pavlidis et al. 2010). To summarize genetic variation of observed and simulated data we calculated multiple population genetics summary statistics, including mean and variance across loci: (1) the number of segregating sites, both per population and summed across populations; (2) nucleotide diversity (π), both per population and for all populations combined; (3) Watterson’s theta per population and for all populations combined; (4) pairwise F_ST_ between populations; the number of shared alleles between pairs of populations; (5) the number of private alleles per population and between pairs of populations; and (6) the number of fixed alleles per population and between pairs of populations. These summary statistics were used as feature vectors on a neural network (nnet) approach implemented in Keras designed to estimate the classification probability of the three simulated models given our data and associated demographic parameters.

After careful parameter exploration, the final architecture of our neural network had three hidden layers with 32 internal nodes and a “relu” activation function. For model classification, our output layer was composed of three nodes and a “softmax” activation function. Three-quarters of the simulations were used as training data, and the remaining 25% were used to test the accuracy of our approach in assigning simulations to the correct model. The neural network training step was run for 1,000 epochs using the “adam” optimizer and a batch size of 20,000 using 5% of the data for validation. To track improvements in model classification during training we calculated the overall accuracy and the sparse_categorical_crossentropy for each epoch. After identifying the most probable model for our observed data, we estimated demographic parameters with a neural network designed to predict continuous variables. Here, we used a similar architecture to the one described above but set an output layer with a single node and a “relu” activation. We used this approach to estimate the effective size of each population, the amount of gene flow between populations, and divergence times. To assess improvements in accuracy during training, we used the mean absolute percentage error (MAE) as an optimizer. We trained the neural network for 3,000 epochs with a batch size of 10,000 and a validation split of 0.1. To account for variation in parameter estimation, we ran 10 replicates and summarized the results calculating mean values for each demographic parameter. We also assessed the accuracy of parameter estimation, by calculating the coefficient of correlation between estimated and true simulated values of the testing data set.

### Testing the link between chromosome length, linked selection, and introgression

To test for predicted patterns associated with linked selection across the genome, we calculated nucleotide diversity (π), relative divergence (F_ST_), and absolute divergence (d_xy_) across the genome. To calculate [, d_xy,_ and F_ST_, we used pixy, a command line utility that handles missing data by adjusting the site-level denominators and sequence length (Korunes and Samuk 2021). Summary statistics were estimated based on vcf files with invariant sites included. To examine the robustness of these estimates given the incomplete nature of GBS datasets, we employed pixy for multiple sliding window widths and data filters (See Supplemental Materials). We specifically examined pixy results for 1) 50 kb windows with an unfiltered vcf; 2) 50 kb windows with a vcf filtered for sites with a minimum depth of 10X, and 3) 250 kb windows with a vcf filtered for sites with a minimum depth of 10X. As we found that pixy results were robust to these depth filters and window sizes (see Supplemental Materials), we opted to use the dataset that retained the most variants, the 50 kb windows with minimum depth of 6X.

To better understand how topology and rates of gene flow vary among chromosomes, we fit data to generalized linear models (glms) in R (R Core Team 2019) to test for an association with chromosome length and bootstrapped these models to examine their sensitivity (see Supplemental Materials). In so doing, we make the assumption that genome structure is relatively conserved across New World suboscine birds (Passeriformes, Tyrannides), since the chromosome lengths of the reference genome are derived from those in *Chiroxiphia lanceolata*, a manakin. This assumption is supported given the high synteny within passerine birds (Dawson et al. 2007; Ellegren 2010; Delmore et al. 2018; Coelho et al. 2019; Peñalba et al. 2020). We first modeled π, d_xy,_ and F_ST_ (unfiltered 50kb-window dataset) as functions of chromosome length (glm family=“gaussian”) based on the results from the pixy analyses outlined above. To do this, we calculated the average value for each statistic across each chromosome by taking the mean value across all windows after eliminating empty windows (i.e., NA values). As the distribution of chromosome lengths was right-skewed, we evaluated these models after log_10_ transforming the data. This allowed us to track general patterns associated with linked selection and recombination in the data. For example, since prior studies have shown that gene flow is expected to increase on smaller chromosomes (Martin et al. 2019), understanding how patterns of divergence, genealogy, and of introgression change with chromosome length enables us to evaluate whether evidence for certain phylogenetic topologies might be driven by gene flow between non-sister populations. For all gaussian regressions, we also estimated Pearson’s correlation coefficient (*r*).

Next, we used logistic regression (family=”binomial”) to test for an association between chromosome length and phylogenetic topology, scoring genetrees (i.e., RAxML trees from the sliding window analysis above) as monophyletic (1) or non-monophyletic (0) for populations in Rondônia, Inambari, and S Rondônia+E Inambari. These logistic regression analyses enabled us to test for associations between genealogical relationships across the genome and chromosome length. ASTRAL recovered two alternative topologies across the genome corresponding to non-monophyletic (T1) and monophyletic (T2) Rondônia populations, and mtDNA recovered a sister-relationship between E Inambari and S Rondônia (T3). Thus, these analyses were aimed at identifying how support for these three topologies varied across the genome to better evaluate whether certain topologies are associated with introgression. For logistic regression, we estimated the coefficient (*β*) and odds-ratio (*ψ*) for each model. We ran these logistic regression models on all 4,858 genetrees as well as a subset of 1,727 genetrees reconstructed from windows with a minimum of 10 SNPs. The latter allowed us to examine the robustness of these models to potential gene tree error. We also replicated this analysis on the chromosome-level ASTRAL phylogenies instead of genetrees.

To obtain direct measures of introgression, we then estimated the introgression proportion (*f*_dM_) across the genome using window-based ABBA-BABA tests in non-overlapping windows with exactly 100 SNPs using ABBABABAwindows.py (https://github.com/simonhmartin/genomics_general (Martin et al. 2014). An initial analysis using windows of fixed width (50 kb and 250 kb) produced highly biased *f*_dM_ estimates, with high variance in *f*_dM_ estimates for larger chromosomes. This was due to the incomplete nature of GBS datasets and amplification bias that led to denser sequence coverage on smaller chromosomes and sparser sequence coverage on larger chromosomes (DaCosta and Sorenson 2014) (see Supplemental Materials). Thus, we chose to define windows by the number of SNPs to help reduce this bias and generate windows with equivalent information content. We specifically explored introgression between N and S Rondônia, which were defined by positive values of *f*_dM_ assuming T1 as the species tree (P1=WI, P2=SR, P3=NR, out=reference; see Results).

These introgression results were then used to model *f*_dM_ and the proportion of derived alleles shared by N and S Rondônia per window (variants shared by P2 and P3 but not P1 or P4; i.e., ABBA’s) as functions of chromosome length to understand how introgression varied across the genome. As positive values of *f*_dM_ indicate introgression between P2 and P3 (N and S Rondônia), we modeled positive *f*_dM_ values as a function of chromosome length after excluding negative values from the data frame. To evaluate whether introgression on the Z-chromosome was lower than mean autosomal introgression, we also used an unpaired Bayesian *t-*test for distributions with unequal variances implemented in the R-package Bolstad (Curran 2013) to statistically test differences in mean values of *f*_dM_ (p2<->p3) across windows on the Z-chromosome versus the mean *f*_dM_ (p2<->p3) for windows on all autosomes combined.

Finally, previous work has shown that recombination not only increases on small chromosomes, but also on the periphery of large chromosomes (Haenel et al. 2018). To test for a potential effect of increased gene flow on the periphery of large chromosomes, we modeled *f*_dM_ (p2<->p3) as a function of percent distance from a chromosome’s center (defined as the distance in base pairs of a window from the center of a chromosome divided by half the chromosome’s length in base pairs). All statistical analyses were performed in R version 4.1.2 (R Core Team 2019). From all analyses, we excluded the W-chromosome, which does not undergo crossing-over and is subject to error given many males in the dataset. We also removed Chromosomes 31, 32, and 33, for which we recovered relatively little data.

## Results

### GBS assembly

Our final GBS assembly included a total of 118,127 loci with a mean and standard deviation of 62,882.65 and 17,506.31 loci assembled per sample, respectively (range=11,955– 85,123 loci). Within this dataset, we recovered a total of 1,277,143 single nucleotide polymorphisms (SNPs).

### Population structure and ancestry

Our results point to between two and four populations across the region corresponding to W Inambari, E Inambari, S Rondônia, and N Rondônia, including individuals with a shared coefficient of ancestrality between W Inambari and S Rondônia, as well as between N and S Rondônia (Fig. 2). Although the best-fit value for the number of ancestries was *k*=2 (Fig. S1), corresponding to populations east and west of the Madeira River, higher values of *k* were consistent with the PCA results, recovering additional distinct ancestries across the Aripuanã (*k=3*) and Purus (*k*=4) Rivers. Contrastingly, PCA consistently recovered four clusters of samples corresponding to areas delimited by the three focal rivers (Fig. 2). These PCA clusters match the spatial boundaries of mitochondrial clades from a previous study (Thom and Aleixo 2015). Still, in consistence with the sNMF results, PC1 (explaining 8.5% of the genomic variation) separated populations from across the Madeira River, PC2 (explaining 6.8% of the genomic variation) separated populations across the Aripuanã River, and PC3 (explaining 2.9% of the genomic variation) separated populations across the Purus River.

**Figure 2:**
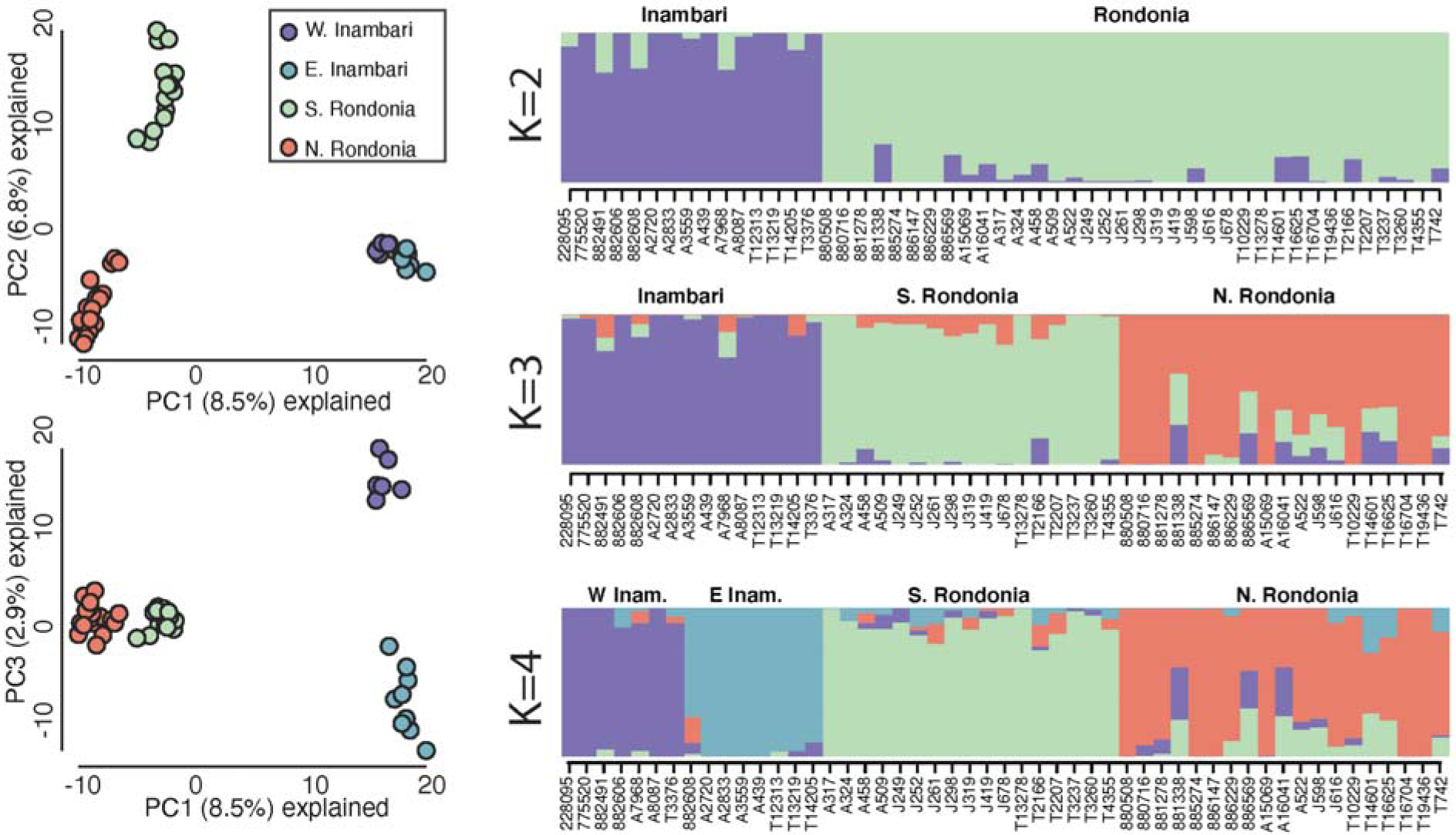
Results of PCA and ancestry on the full GBS dataset from samples spanning four populations. Colors correspond to shaded regions on the map (Fig. 1): W Inambari (purple), E Inambari (teal), S Rondônia (pale green), and N Rondônia (red). (Left) Results of PCA on all samples. (Right) sNMF results for the optimal K-value of K=2 (top), as well as plots for K=3 (center) and K=4 (bottom).

### Genetrees, chromosome trees, species tree, and demographic history

We detected three competing phylogenetic topologies across the genome, two of which predominated among chromosome trees (Fig. 3): Topology 1 (T1) matched the genome-wide species tree and was recovered for nine of the 34 chromosomes, including the Z-chromosome; T2 matched the autosomal tree and was recovered for 22 of the autosomes. There were also three unique topologies recovered for Chromosome 29, Chromosome 31, and Chromosome 32 (Fig. S2). None of these were concordant with the mitochondrial topology recovered in a previous study (henceforth, T3) (Thom and Aleixo 2015). Overall, populations from Rondônia were recovered as monophyletic for 22 chromosome trees and Inambari populations were monophyletic for all but three chromosomes.

**Figure 3:**
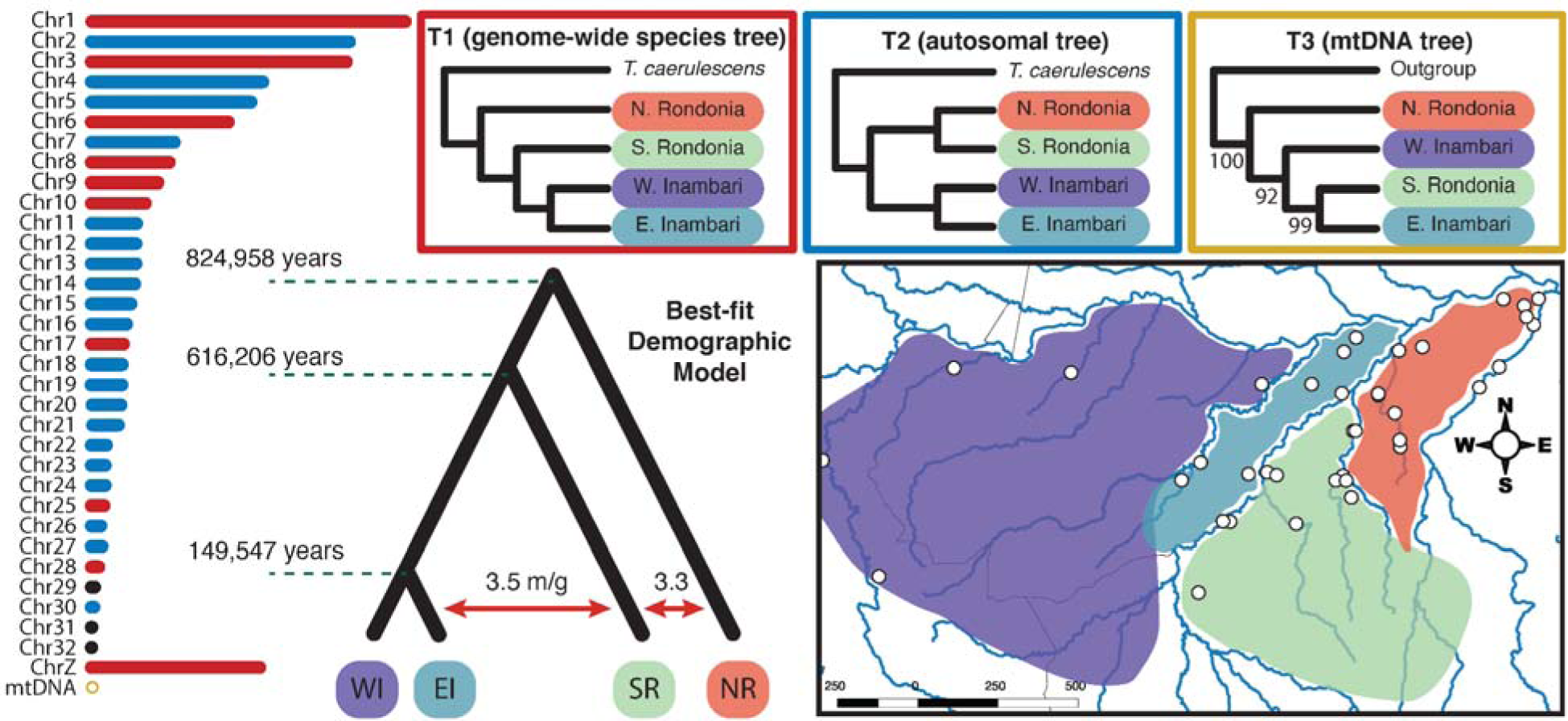
Chromosome-level topologies inferred in astral for 50 kb sliding windows across the genome. Two chromosome tree topologies predominated across the genome: T1 (red chromosomes/red box), which is consistent with the genome-wide tree, and T2 (blue chromosomes/blue box), which is consistent with the autosomal tree. The mitochondrial topology is also shown (T3) with bootstrap support, based on the results of a previous study (Thom and Aleixo 2015). The tree at the lower left shows the best-fit demographic model inferred using a convolutional neural network. Bars in the left column are proportional to chromosome length.

Demographic modeling similarly recovered T1 as the population history (model classification probability: model 2 = 0.99), finding high gene flow between S Rondônia and E Inambari as well as between N and S Rondônia (Fig. 3). Model classification yielded high accuracy, with training data being correctly assigned to the simulated model 99% of the time (accuracy = 0.99). The neural network regression approach designed for demographic parameter estimations produced accurate estimates for most parameters (R^2^ simulated ∼ estimated > 0.90), except for the two oldest divergence times, likely due to the high amount of gene flow between populations (R2 ∼ 0.61; Table S4). Effective population sizes were in general consistent with the size of the geographic distribution of the species, with the most restricted E Inambari populations having the smallest size (198,314 individuals; R^2^ = 0.93; MAE = 72,272). The divergence between E and W Inambari populations occurred around 150 Ka (149,546ya; R^2^ = 0.95; MAE = 48,492), followed by the split between the ancestor of the Inambari populations from S Rondônia at about 600 Ka (616,206ya; R^2^ = 0.61; MAE = 117,941), and the deepest divergence at around 820 Ka (824,957ya; R^2^ = 0.61; MAE = 125,875). Gene flow estimates were high between both migration edges of the model suggesting considerable introgression between E Inambari and S Rondônia, and between Rondônia populations. Parameter estimates were in general contained within simulated priors except for gene flow estimates. Additional runs adjusting priors for gene flow drastically affected the accuracy of model classification, thus we assumed these constrained and conservative estimates.

The support for the three alternative topologies varied across the genome (Fig. S3 and S4; Table 1). The unrooted T1/T2 topology (T1 and T2 are the same when the ingroup quartet is unrooted) had the highest support across the genome in our analysis for the ingroup populations. However, when including the reference to differentiate between T1 and T2 topologies, both T1/T3 (T1 and T3 are the same when this quartet is unrooted) and T2 had similar support across the genome. There were relatively few outlier loci supporting T3 compared with the other topologies, and we did not find any association between n-mt genes and outlier peaks supporting the mitochondrial topology.

**Table 1:** Number of outlier windows supporting three alternative topologies (T1/T2, T3, and a third unrooted topology Tx), and the overall number of genes, and number genes linked to mitochondrial activity less than 100kb from outlier peaks (N-mt genes) on figure S4 [ingroup TWISST analysis].

| Unrooted Topology | Number of outlier windows | Number of genes | Number of N-mt genes | Number of windows with highest weight |
| --- | --- | --- | --- | --- |
| T1/T2 | 43 | 262 | 19 | 1586 |
| Tx | 8 | 26 | 1 | 360 |
| T3 | 10 | 17 | 1 | 441 |

### Associations with chromosome length

We detected significant associations between chromosome length, genetree topologies, and genome statistics (Fig. 4). First, we found a significant association with F_ST_ (*n*=32, 50kb windows: *r*=0.818, *P*<0.0001; 250kb windows: *n*=32, *r*=0.666, *P*<0.0001; Fig. 4A), d_xy_ (50kb windows: *n*=32, *r*=-0.856, *P*<0.0001; 250kb windows: *n*=32, *r*=-0.931, *P*<0.0001; Fig 4B), and π (S Rondônia: 50kb windows, *n*=32, *r*=-0.885, *P*<0.0001, 250kb windows, *n*=34, *r*=-0.868, *P*<0.0001; 50kb windows N Rondônia: *n*=32, *r*=-0.883, *P*<0.0001; 250kb windows: *n*=34, *r*=-0.846, *P*<0.0001; Fig. 4C and 4D). Bootstrapping (Fig. S5) and pairwise correlations based on alternate VCF filtering and window-sizes showed that the results of our linear regression models were not affected by unequal sequence coverage across windows and chromosomes (Fig. 5, S6, and S7). We recovered a negative association between chromosome length and genetree topology, wherein genetrees on larger chromosomes have a reduced probability of recovering monophyly of Rondônia (*β*=-0.231, *ψ*=0.794, *P*=0.0005; Fig. 4F) and recapitulated this result at the level of chromosomes, though the latter was only weakly supported (*β*=-0.0003, *ψ*=0.999, *P*=0.054; Fig. 4E).

**Figure 4:**
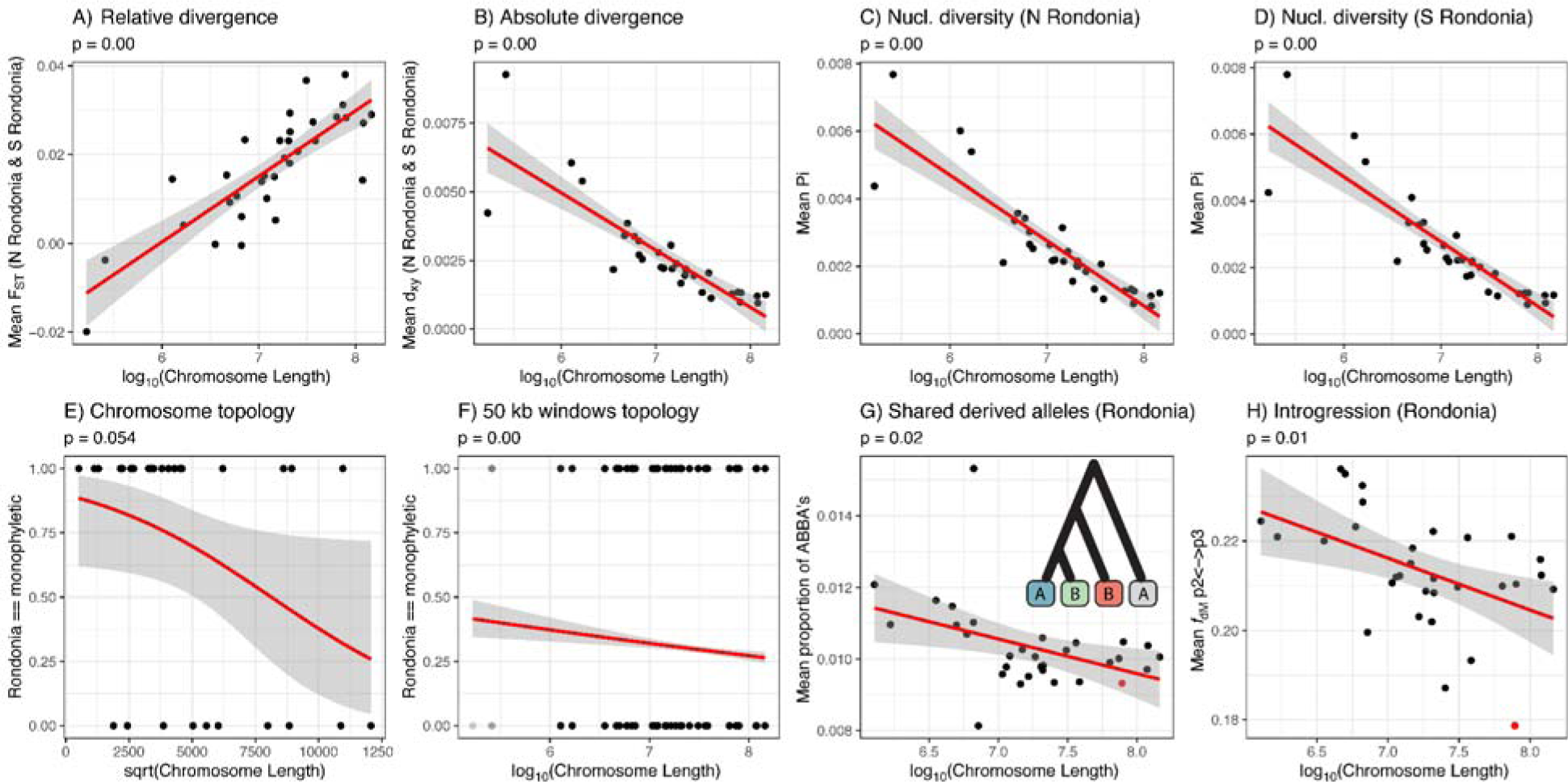
Logistic and linear regression tests for associations between genomic characteristics and chromosome length. Y-axis values in the top row reflect chromosome-wide means of 50-kb windows for (A) F_ST_, (B) d_xy_, (C) pi for N Rondônia, and (D) pi for S Rondônia. Plots in the bottom row examine the influence of introgression on genetree variation across the genome showing (E) a nonsignificant association between chromosome topologies and chromosome length despite (F) a significant negative association in genetree topologies consistent with T2 and chromosome size, and (G and H) significant negative associations between introgression and chromosome length. Models for each association are shown as the red line with the shaded gray area representing the model standard error. Red points in plots G and H represent values for the Z-chromosome. Derived variants shared by Rondônia were defined as SNPs showing an ABBA pattern assuming topology T1, as indicated using the tree at the top right of plot G. The topology tips, from left to right are E Inambari, S Rondônia, N Rondônia, and *T. caerulescens* (outgroup).

**Figure 5:**
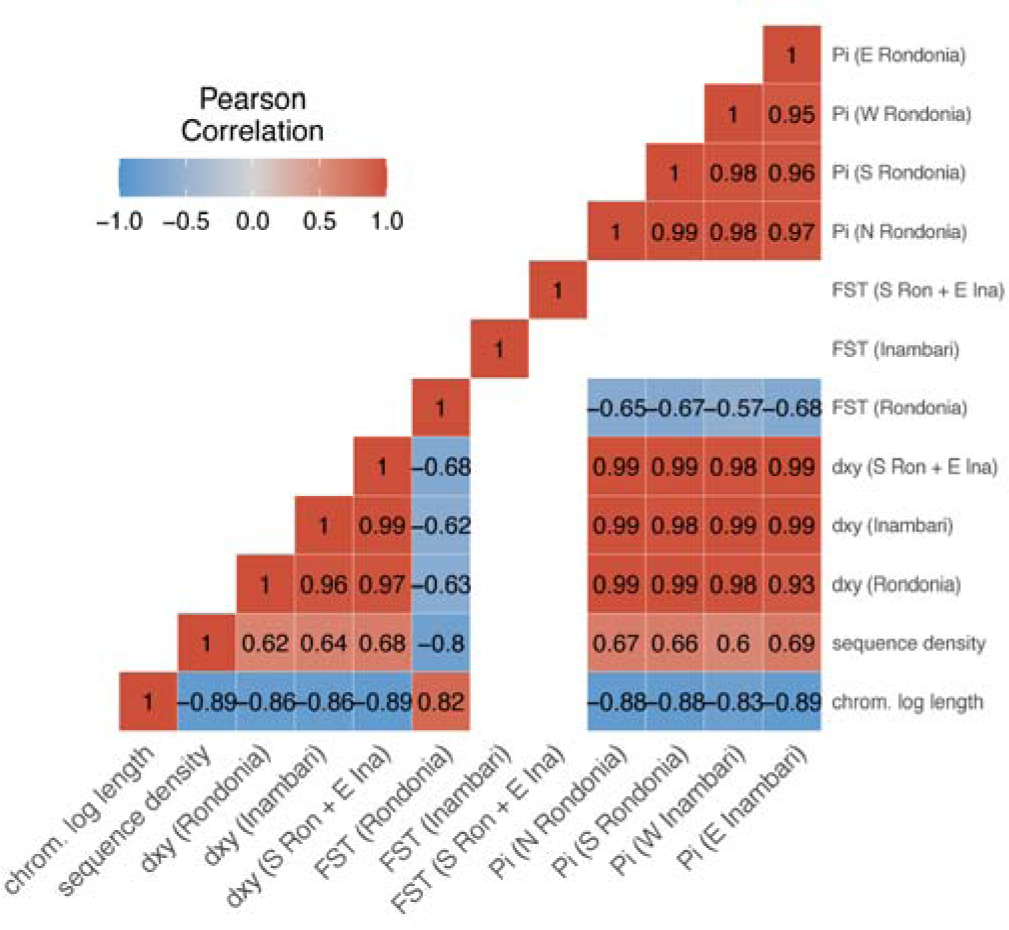
Heatmap of Pearson’s coefficients for pairwise correlations showing pairwise correlations of within-chromosome means of genetic summary statistics for values of d_xy_, F_ST_, and pi. Empty (white) boxes represent correlations with nonsignificant P-values (*P*>0.05).

We also confirmed that introgression varied across the genome and was negatively correlated with chromosome length (Fig. 4G and 4H). Specifically, we found significant negative correlations between *f*_dM_ (*n*=31, *r*=-0.476, *P*=0.007) and the proportion of derived variants shared by N and S Rondônia (ABBA’s; n=31, *r*=-0.420, p=0.019) with chromosome length. The Bayesian t-test confirmed (*df*=237.74, *t*=3.566, *P*=0.0005) that the Z-chromosome had lower levels of introgression (mean *f*_dM_=0.178) than autosomes (mean *f*_dM_=0.212). Overall, *f*_dM_ values averaged positive across the genome, implying stronger gene flow between N and S Rondônia than between N Rondônia and E Inambari (positive values indicate an excess of gene flow between P2 and P3, whereas negative values indicate an excess between P1 and P3). We found no relationship between *f*_dM_ and distance from chromosome center (Fig. S9).

## Discussion

### Nonrandom variation in genealogical history across the genome

We combine phylogenomics and population genetics to investigate the interplay between genomic architecture and biogeographic processes in generating predictable patterns of genetree variation across the genome of an Amazonian antbird, *Thamnophilus aethiops*. We found that accounting for chromosome length can inform phylogenetic and biogeographic inference in cases of high gene flow among non-sister taxa. Our results also suggest that reduced representation genomic data such as genotype-by-sequencing can be used with genomic-architecture-aware approaches, recapitulating expected associations between genomic processes and the signal for ancestry and introgression.

We tested three competing hypotheses for the relationships of four spatially adjacent and genetically differentiated populations that are semi-isolated across Amazonian tributaries. Two of these topologies, T1 and T2, were equally supported across genome-wide sliding windows (Fig. S3), which made inferring *T. aethiops*’ evolutionary history challenging. We found that genealogical signal for these competing hypotheses was nonrandomly distributed across the genome; areas of low gene flow such as the Z-chromosome and larger autosomes tend to support T1, whereas areas with elevated gene flow such as smaller autosomes tend to recover genetrees consistent with T2 (Fig. 4E–F). The third topology, T3, was not well supported across the nuclear genome (Fig. S4), but was recovered for mitochondrial DNA. Importantly, introgression was negatively correlated with chromosome length and the Z-chromosome exhibited especially lower introgression than autosomes, consistent with expectations (Fig. S10). Thus, we suggest that T1 may be seen as representing the initial branching pattern among the four taxa, whereas the prevalence of T2 on smaller autosomes probably resulted from introgression between N and S Rondônia. These results suggest that the genome-wide diversification history of *T. aethiops* might be better explained by a complex network of differentiation and introgression between multiple interacting populations.

The idea that the prevalence of T2 is driven by gene flow is supported by theoretical predictions about genome architecture. Smaller chromosomes are expected to exhibit higher levels of gene flow than larger chromosomes due to their higher recombination rates which more effectively break the linkage between introgressed variants (Martin et al. 2019; Tigano et al. 2022). During Meiosis, each chromosome must undergo at least one crossing-over event resulting in fewer recombination events per base on longer chromosomes than smaller ones. This becomes complicated in birds, which may also experience increased selection on smaller chromosomes due to denser gene content in those regions (Henderson and Brelsford 2020). However, in birds, recombination is also thought to be tied to gene promoter regions (Singhal et al. 2015). Thus, although there is an increase in the density of targets of selection along smaller chromosomes, there is also additional recombination acting to reduce the effects of such linked selection. To test this assumption, we modeled multiple population genetic statistics as functions of chromosome length. We found that F_ST_ decreased on smaller chromosomes, but π and d_xy_ were negatively correlated with chromosome length. These patterns are consistent with expectations associated with a reduction in overall genetic diversity due to linked selection on larger chromosomes, thereby increasing F_ST_ and decreasing d_xy_. We also found that introgression statistics such as *f*_dM_ and the proportion of derived alleles shared by Rondônia populations (i.e., ABBA’s given T1) were negatively correlated with chromosome length. Overall, these results confirm our assumption that rates of gene flow between N and S Rondônia are higher on smaller chromosomes and likely driven by increased recombination.

### Biogeographic and geogenomic implications

Understanding how variation in genealogical history is associated with genomic processes enables a closer look at the processes driving population divergence and speciation in *T. aethiops,* and can illuminate our understanding of Amazonian biogeography in general. Amazonian biogeography has been at the center of discussions on how landscape evolution leads to allopatric speciation (Haffer 1997; Ribas et al. 2012), but increasingly, researchers are discovering that the histories of taxa across this landscape are marked by high gene flow (Barrera-Guzmán et al. 2022). Recent genomic studies have reported introgression across rivers and considerable phylogenetic conflict, often despite strong genetic and phenotypic structuring (Thom et al. 2021; Del-Rio et al. 2022; Musher et al. 2022). This includes examples of hybrid speciation (Barrera-Guzmán et al. 2018), mitonuclear discordance (Del-Rio et al. 2022), mitochondrial capture (Ferreira et al. 2018), and extensive introgression across river headwaters (Weir et al. 2015). Phylogenetic relationships among interfluvial populations have been used to inform paleogeographic models of landscape evolution and have also helped to generate and test multiple biogeographic hypotheses (Cracraft and Prum 1988; Ribas et al. 2012). However, the unique configuration of the Amazon Basin with massive, unstable tributaries flowing in parallel facilitates episodic or continuous gene flow, which can result in reticulate patterns of differentiation (Barrera-Guzmán et al. 2018; Thom et al. 2018). This process directly affects phylogenetic inference and, if not fully understood, hampers researchers from obtaining a detailed understanding of the region’s biogeographic history. Our study highlights how genealogical patterns vary predictably across the genome and inform biogeographic inference (Martin et al. 2019).

We suggest that large-scale river capture events can result in historical signatures of discordant genealogy across the genomes of species that respond to rivers as barriers (Musher et al. 2022). The river capture scenario postulated in prior studies (Fernandes 2013; Weeks et al. 2016; Ruokolainen et al. 2019) predicts a sister relationship between Inambari and S Rondônia with gene flow between E and W Inambari as well as between N and S Rondônia. Thus, either T1 or T3 might match the spatial-phylogenetic expectations under a river-vicariance model. However, our results are consistent with a more nuanced set of expectations associated with barrier change wherein genomic heterogeneity is associated with multiple distinct genealogical histories. Assuming that T1 reflects the history of population isolation (or at least reduced introgression) across the Aripuanã, then T2 results from secondary contact and lineage fusion within the traditionally recognized area of endemism, Rondônia (Cracraft 1985). This gene flow among N and S Rondônia appears to be resulting in autosomal homogenization; sNMF preferred a model of *k*=2, recovering two ancestral populations corresponding to Inambari and Rondônia (Fig. 2 and S1). Within Rondônia, the lack of any apparent plumage variation in *T. aethiops* also supports the notion that there is homogenizing gene flow between the N and S Rondônia populations.

A key objective of this study was to dissect the interplay between genomic and biogeographic processes in generating genomic heterogeneity, which requires some knowledge about landscape history. Recently, researchers have proposed a field of study, called “geogenomics,” wherein patterns of genomic differentiation and genomically-inferred timings of divergence and gene flow can be used to help test paleogeographic models (Dawson et al. 2022; Ribas et al. 2022). As shown here and elsewhere, spatial diversification patterns within Amazonia are reticulated, and unraveling the evolutionary history of taxa in this system is nontrivial (Dagosta and Pinna 2017; Dagosta and De Pinna 2019). Understanding the geological context of river capture while accounting for intrinsic genomic processes, however, aids in the interpretation of alternative phylogenetic histories. For instance, if the Madeira headwaters were captured within the past few hundred thousand years, the Aripuanã River must predate the upper Madeira, as implied by both T1 and T3. Given that T1 is probably less impacted by introgression and populations within Rondônia are now homogenizing, the Aripuanã probably represented a more important barrier for *T. aethiops* prior to the capture event that is now weakened. In this way, we have a window into the process driving the formation of areas of endemism. As new barriers form on the landscape, old barriers erode; if differentiated taxa are not reproductively isolated, as appears the case for *T. aethiops* populations in Rondônia, they may fuse into a single taxon whose distribution conforms to the boundaries of the new river-barriers, leaving behind only reciprocally monophyletic mitochondrial groups that potentially match the ancestral landscape configuration. Extensive paleochannels between the Jiparaná and Aripuanã tributaries of the middle Madeira Basin suggest that these rivers have historically been larger and behaved as a dynamic megafan, a hypothesis also supported by biological data (Latrubesse 2002; Wilkinson et al. 2010; Ferreira et al. 2017). We thus postulate a historical river somewhere in the vicinity of these two tributaries that could have acted as a historical barrier to dispersal. Under this scenario, the paleo-Madeira River would have been flowing via the Jiparaná or Aripuanã basins or somewhere in between (Hayakawa and Rossetti 2015), acting as a historical barrier for taxa located on either side. Likewise, the Tapajós basin to the east was probably drained via this paleo-Madeira River (Rossetti 2014). If so, the formation of the modern Tapajós would have drawn water away from this basin, reducing the barrier’s strength. Once the Purus tributary was captured, the modern Madeira formed by extending its headwaters, which in turn generated a new barrier on the landscape. If taken at face value, our best-fit demographic model suggests that this river capture occurred roughly 600 kya, at least 400 ky prior to the current geological estimate (Ruokolainen et al. 2019), but in line with divergence times of some other taxa in the region (Silva et al. 2019; Musher et al. 2022).

### Interpreting the mitochondrial topology

Given that T1 and T2 are the primary genealogical signals across the genome, we are left to evaluate the mechanisms that gave rise to the mitochondrial tree (T3). The dispersal ecology of resident Amazonian bird species is poorly understood, but male birds are generally considered philopatric, which means it is unlikely that male-biased dispersal drives mtDNA patterns. T3 was estimated based on two loci, cytochrome-*b* and NADH-dehydrogenase subunit 2, and is well-supported (Thom and Aleixo 2015). Mitochondrial DNA is known to have faster coalescent times due to its reduced (one quarter) effective population size and has thus traditionally been viewed as an efficacious phylogenetic marker for detecting divergences that occur over short timescales (Zink and Barrowclough 2008). However, mtDNA is a single locus and therefore might be expected to disagree with the species tree under certain conditions (Maddison 1997). For example, if we assume T1 to be the “true” species tree, then T3 may have resulted from mitochondrial capture during hybridization between E Inambari and S Rondônia, as documented in other groups (Ferreira et al. 2018; Myers et al. 2022). If this were the case, we should find portions of the genome that are associated with the cell respiratory system to be introgressed due to the same event (Morales et al. 2018). Yet, there is limited support for T3 across the genome (Fig. S4), and the windows that do support T3 are not clearly linked to genes associated with mitochondrial activity (Table 1). However, because we used GBS data for this analysis, we could be missing crucial mitonuclear gene clusters in our dataset.

Alternatively, incomplete lineage sorting or biogeographic history could have given rise to the mtDNA topology. For example, it could have resulted from a deep coalescent event wherein W Inambari haplotypes failed to coalesce with E inambari haplotypes. This is especially likely in instances of very rapid divergence (Degnan and Rosenberg 2006). Given that divergences in our ingroup occurred in under one million years, this is certainly plausible, despite the rapid fixation rate of mtDNA. However, it is also possible that T3 represents the history of population isolation across rivers (Fig. 1); it is striking that the distribution of reciprocally monophyletic populations based on mtDNA (Thom and Aleixo 2015) matches the spatial patterns recovered in the PCA and that *T. a. injunctus* (E Inambari) is most phenotypically similar to *T. a. punctuliger* (Rondônia). Therefore, if the T3 topology indicates the signal of population isolation across rivers then, introgression between multiple non-sister lineages has nearly erased the signal of that isolation from the genome since relatively few nuclear loci support T3 (Table 1). In this case, T1 does not exactly show the history of isolation across rivers, but might just reflect fewer genetic incompatibilities between E and W Inambari, which are more recently diverged, than between N and S Rondônia. In other words, T1 may not reflect a lack of gene flow altogether, but instead easier gene flow between E and W Inambari, which lack as many incompatibilities.

The genomes of populations that diversify on dynamic landscapes contain the signatures of multiple histories; that is, they are reticulated. To dissect these histories of isolation and secondary contact, we argue that it is important to understand the biogeographic mechanisms that give rise to predictable genomic patterns. If one‘s objective is to model the relationships of taxa –i.e., reconstruct phylogeny– T1 is the best supported topology given our phylogenetic and demographic modeling results. However, if the goal is to decipher the complex biogeographic history of these taxa across space and time, we may want to know whether the mtDNA tree resulted from mitochondrial capture, deep coalescence, historical isolation, or even simply phylogenetic error. We suggest that T3 could represent the historical population divergences across rivers as they formed. The divergence time based on mtDNA between E Inambari and S Rondônia occurred at roughly 200 kya (Thom and Aleixo 2015), which lines up well with geological estimates for the Madeira River capture event (Ruokolainen et al. 2019). However, future studies based on whole-genome resequencing data may be necessary to fully understand the complicated patterns of isolation, reticulation, and homogenization in *T. aethiops*, and the relationships between these processes and genome architecture. It would also enable more detailed models of historical demography and selection not possible with GBS data.

### Concluding remarks

Gene flow may be a creative or destructive force with regards to divergence and speciation. Introgression among divergent populations, where standing genetic variation among populations is episodically reshuffled into novel combinations might be an underappreciated speciation mechanism (Marques et al. 2019). We showed that introgression remains high among taxa of a common understory antbird, *T. aethiops*. Specifically, taxa within *T. aethiops* seem to remain differentiated, despite ongoing and apparently intense introgression. This has been shown in many other Neotropical taxa (Martin et al. 2013; Ebersbach et al. 2020; Musher et al. 2022). Given the potential for high rates of episodic isolation and reconnection due to the movement of large rivers, conditions in Amazonian lowlands, like those in some African lakes (Aguilée et al. 2013; Meier et al. 2017), are potentially ideal for this combinatorial mechanism of adaptation to promote diversification in lowland birds. Still, the process of high introgression can also result in higher rates of extinction (i.e., homogenization) for these young, weakly differentiated taxa, as they fuse with other lineages (Harvey et al. 2017; Barrera-Guzmán et al. 2022). Nonetheless, our results suggest that a nontrivial portion of genealogical heterogeneity across the genome arises due to extrinsic processes –such as river-course rearrangement– interacting with intrinsic processes associated with genome architecture.

## Acknowledgments

We are grateful to E. Griffith, A. Del Grosso, J. Weckstein, and J. Merwin for comments on an earlier draft of this manuscript. We also thank B. Faircloth for advice and assistance, as well as the Bauer Core Facility and Cannon/Odyssey HPC system staff at the Harvard University Faculty of Arts and Sciences for support and computational resources needed for sequencing and assembly of the *T. caerulescens* reference genome. We thank S. V. Edwards and the Department of Organismic and Evolutionary Biology at Harvard University for providing financial support to generate the reference genome of *Thamnophilus caerulescens*. LJM was funded in part by NSF grant DEB-1855812 to J. D. Weckstein. We also thank curators and staff at the Instituto Nacional de Pesquisas Amazonica, Museu Paraense Emilio Goeldi, American Museum of Natural History, and Louisiana Museum of Natural Science for loaning invaluable genetic resources. Reviews from F. Burbrink and two anonymous referees greatly improved the quality of this manuscript.

## Data availability

Supplemental material and scripts are available https://datadryad.org/stash/share/KB1ZlWN6mIs_FJvI7OWpb8wqQiJT50OWQPVpITQakSs

