## Supplemental Information for "Geogenomic predictors of genetree heterogeneity in an Amazonian bird (*Thamnophilus aethiops*)"

**Supplementary Materials for “Geo-genomic predictors of genetree heterogeneity in an Amazonian bird (*Thamnophilus aethiops*)*”***

**Glaucia Del-Rio,** Cornell Laboratory of Ornithology and Department of Ecology and Evolutionary Biology, Cornell University, Ithaca, NY, USA

**Rafael S. Marcondes,** Department of Biology and Museum of Natural Science, Louisiana State University; Department of BioSciences, Rice University, USA

**Robb T. Brumfield,** Museum of Natural Science and Department of Biological Sciences, Louisiana State University, Baton Rouge, LA 70803, USA

**Gustavo A. Bravo,** Sección de Ornitología, Colecciones Biológicas, Instituto de Investigación de Recursos Biológicos Alexander von Humboldt, Claustro de San Agustín, Villa de Leyva, Boyacá, Colombia; Museum of Comparative Zoology and Department of Organismic and Evolutionary Biology, Harvard University, Cambridge, MA, USA

**Gregory Thom,** Museum of Natural Science and Department of Biological Sciences, Louisiana State University, Baton Rouge, LA 70803, USA

**INTRODUCTION**

Herein, we present additional analyses, results, and discussion on genomic patterns in *T. aethiops.*

**METHODS**

*Reference genome sequencing and assembly*

The *T. caerulescens* reference genome was generated based on a female specimen collected in 2018 in Bolivia (Departamento Cochabamba: Siberia) and deposited at the Louisiana State University Museum of Natural Science (tissue number B-94639, voucher skin number LSUMZ 227821). We followed the ChromiumTM Genome Protocol for high molecular weight genomic DNA extraction from fresh frozen tissue available at https://support.10xgenomics.com/genome-exome/sample-prep/doc/demonstrated-protocol-hmw-dna-extraction-from-fresh-frozen-tissue. Using the Genomic DNA protocol of the Agilent 4200 Tape Station System, we verified that the DNA integrity index was 9.3 and that the percentage of fragments > 50k bp was greater than 50%. Library preparation and sequencing for 10X Genomics linked-reads was conducted at the Bauer Core Facility of the Harvard University FAS Division of Science. The paired-end library was sequenced on a single lane of a NovaSeq SP 2 x150bp. We used Supernova v 2.1.1 (Zheng et al. 2016; Marks et al. 2019) to demultiplex, assemble, and generate fasta files. This initial assembly was conducted using all reads (829.66 M), which yielded an effective coverage of 68X, contig N50 of 166.66 Kb, scaffold N50 of 2.97 Mb, and an estimated genome size of 1.24 Gb.

We filtered scaffolds < 1000 bp from the assembly using Kent tools (http://hgdownload.soe.ucsc.edu/downloads.html#source_downloads), and used 10X Genomics Long Ranger software v 2.2.2 (Zheng et al. 2016) to remove the proprietary 10X barcodes from the raw sequencing reads. Then, we mapped the trimmed reads back to the filtered assembly using samtools v 0.1.19 and bwa v 0.7.17 (Li and Durbin 2009; Li 2011). After mapping, we polished the filtered scaffolds using Pilon v 1.23 (Walker et al. 2014), and we used custom Python code to rename scaffolds and convert gaps longer than 50 bp to 100 bp. Because polishing can break some scaffolds, we performed another round of scaffold filtering to remove scaffolds < 1000 bp, and we estimated assembly completeness using BUSCO v 4.0.6 (Seppey et al. 2019). This initial BUSCO analysis suggested a moderate level of duplication for some orthologs, and we decided to remove potential haplotigs from the polished assembly using purge_haplotigs v.1.1.1 (Roach et al. 2018). Because repeat regions can confound haplotig identification and removal, we modeled repeats in the polished assembly using RepeatModeler v 2.01 (Smit and Hubley 2008) before identifying repeats using RepeatMasker v 4.1.0 (Smit et al. 2013) and converting the GFF file of repeat output to BED using awk. After repeat modeling, we re-aligned the trimmed, raw reads to the polished assembly using bwa and samtools; generated a histogram of sequence coverage using purge_haplotigs; selected coverage cutoffs of 25X (low), 50X (mid), and 190X (high); and purged putative haplotigs from the assembly by inputting the coverage estimates, coverage cutoffs, and BED-formatted repeat regions to purge_haplotigs. After haplotig purging, we checked assembly completeness again using BUSCO. We performed a final round of reference-based scaffolding using the high-quality Vertebrate Genome Project assembly of *Chiroxiphia lineola* (GCF_009829145.1; divergence time: ~40Ma) (Koepfli et al. 2015; Harvey et al. 2020). Before scaffolding, we used custom Python code to rename the *Chiroxiphia* scaffolds by chromosome number and to remove unplaced scaffolds and contigs from the *Chiroxiphia* assembly - leaving only a file of chromosomes sequences. Then, we masked repeats in our haplotig purged assembly and used ragtag v 1.0.0 (Alonge et al. 2019) to scaffold our assembly against the *Chiroxiphia* chromosomes with default options (except setting gaps to 200 bp). We renamed the pseudochromosomal scaffolds to reflect the reference genome and chromosomes used to scaffold them using custom Python code. We removed all soft-masking from the pseudochromosomal assembly and performed a final round of repeat masking using RepeatMasker and the repeat models created above. We computed contiguity statistics with Quast v 5.0.2 (Mikheenko et al. 2018) and performed a final round of assembly completeness checks with BUSCO.

We annotated the *T. caerulescens* genome by downloading a copy of vertebrate protein sequences from OrthoDB v 10 (Kriventseva et al. 2019), filtering that file to retain only proteins with a verified IUPAC protein alphabet, and predicting genes and proteins using a containerized version (https://github.com/faircloth-lab/singularity/; commit 2650d41) of braker v 2.1.5 (Hoff et al. 2019) with default options. To associate protein sequences with gene metadata (name, function, database number, etc.), we used blastp v 2.6.0 (NCBI Resource Coordinators 2018) to align the augustus.hints.aa file to annotated proteins from both *Chiroxiphia lanceolata* (GCF_009829145.1_bChiLan1.pri_protein.faa) and *Taeniopygia guttata* (GCF_008822105.2_bTaeGut2.pat.W.v2_protein.faa) with default options except that we set -evalue=1e-10, -max_target_seqs=1, and we output a custom table of values (-outfmt 6). Then, we input each custom table of blastp results, the GPFF file corresponding to each protein fasta file, and the augustus.hints.gtf file from braker to a custom computer program (parse_braker_gtf_and_blast.py; https://github.com/faircloth-lab/singularity/tree/main/braker/scripts; commit 456cce4) to add annotation metadata from each GPFF to the attribute field of each GTF file generated for *T. caerulescens* by braker. This produced two sets of updated annotations (GTF format): one file with annotation metadata from *C. lanceolata* and another with annotation metadata from *T. guttata.* Predicted genes in each GTF file without significant blastp hits were not modified, other than to correctly propagate gene_id and transcript_id values across appropriate lines of each GTF file.

*Evaluation of data biases and sensitivity analyses*

Previous studies have shown that amplification bias can occur in restriction site-associated DNA sequencing data that results in higher sequence coverage on smaller chromosomes, which tend to have higher GC content (DaCosta and Sorenson 2014). To evaluate whether the correlation results for F_ST_, d_xy_, and π might be artifacts of unequal sequencing across chromosomes, rather than derived from the genomic architecture of the species, we first identified possible sequencing bias by plotting sequence density as a function of chromosome length. Sequence density was estimated by taking the number of 50kb windows with sequence coverage divided by chromosome length. We then performed a bootstrapping procedure to reduce the effects of unequal sequence densities across chromosomes of different sizes. At each bootstrap replicate, a subset of random windows was chosen across each chromosome and average values of d_xy_, F_ST_, and π were re-estimated for each chromosome. Because there were relatively few windows with coverage on each chromosome in the 200kb window dataset, we subsampled 25 random windows at each iteration when bootstrapping this dataset. Likewise, as there were many windows with coverage on each chromosome in the 5kb window dataset, we subsampled 100 random windows for this dataset. We then calculated the mean p-value and model coefficient across all runs and plotted their distributions.

To examine whether our depth filter of 6X for statistical base calling might affect summary statistic inference, we tested for a differences in pi, F_ST_, and d_xy_ calculated from 6X and 10X datasets using a Wilcoxon rank sum test.

**RESULTS**

We found our results were robust to potential and observed data biases. We found sequence density was negatively correlated with chromosome length, indicating sequenced DNA was overrepresented on smaller chromosomes relative to large ones (*n*=31, *r*=-0.911, *P*<0.0001). Still, our bootstrapping analyses indicated that linear regression models were robust to this bias (Fig. S5). The Wilcoxon test showed that means of pi (S Rondonia: *W*=276220374, *P*=0.099; N Rondonia: *W*=281517502, *P*=0.163), d_xy_ (Rondonia: *W*=8522565124, *P*=1.0), and F_ST_ (Rondonia: *W*=879355984, *P*=1.0) did not statistically differ for datasets filtered for minimum depth of 6X or 10X, which indicates any potentially erroneous base calls in the 6X dataset were few or absent and did not affect summary statistic inference (and thus demographic and other model-based inferences).

We recovered a slight negative association between chromosome length and genetree topology, wherein genetrees on larger chromosomes have a reduced probability of recovering monophyly of Rondônia (*β*=-0.231, *ψ*=0.794, *P*=0.0005) and Inambari (*β*=-0.172, *ψ*=0.842, *P*=0.007), but not a S Rondônia + E Inambari topology (*β*=-0.064, *ψ*=0.938, *P*=0.437). This result held up for Rondonia monophyly even when only including genetrees built from a minimum of 10SNPs (Fig. S8).

**DISCUSSION**

Scenarios of rapid population differentiation coupled with multiple instances of introgression do not fit neatly into a bifurcating phylogenetic model (Mallet et al. 2016). Complex metapopulation dynamics combined with limited geological information make it difficult to establish direct links between landscape evolution and speciation in Amazonia (Smith et al. 2014). Although phylogenetic relationships might have been affected by multiple pulses of introgression, populations bounded by rivers are in general differentiated from one another, which corroborates the strong effect of rivers as barriers, if not the presence of postzygotic incompatibilities (Pulido-Santacruz et al. 2018). Still, the complex patterns of differentiation and introgression that have been reported for Amazonian species suggest that the use of biological data to inform geological models should be approached with caution. Moving forward, the ability to test more detailed models using genomic data will result in increased information about the history of rivers, thus informing geological knowledge.

Nevertheless, there are certain limitations that preclude us from fully understanding how effective rivers are at isolating populations over timescales necessary for speciation. For example, we still have limited information about the timing of riverine changes, the amount of gene flow or divergence that might be caused by a river-course change, or the variation in gene flow between a river’s mouth and headwaters. Moreover, it is unclear whether results based on simplistic models might mislead inferences about species’ “true” histories. We might expect that if episodic river course changes are the primary mechanism driving introgression among taxa, then gene flow should be restricted to discrete periods matching such landscape changes. In theory, this can be tested with demographic models, but in practice this task is nontrivial; methods can be sensitive to data, which often violate key assumptions of the models employed. These models also do not yet account for intrinsic properties of the genome. Although we cannot differentiate gene flow associated with continuous dispersal from pulses caused by physiographic changes, the concordance we found between the Madeira River course change and phylogenetic expectations suggest that historical changes in the landscape were an important mechanism driving population isolation and connectivity (Fig. 8).

*The impact of amplification biases in our dataset*

Some previous studies have shown that certain types of GBS studies can experience amplification biases that result in among-locus variation in genomic coverage and sequencing depth(DaCosta and Sorenson 2014). This in turn results in overrepresentation of sequence coverage in GC-rich portions of the genome. Our results strongly match these predictions as we found similar biases in our own dataset: we detected a strong negative correlation between chromosome length and sequence density (Fig. 5). However, these results seem not to have biased most downstream analyses. For example, our bootstrapping exercise revealed that randomly sampling equivalent numbers of windows from each chromosome recovered similar model results as modeling all windows across all chromosomes (Fig. 7). Likewise, statistically significant values for Pearson’s correlation coefficient (*r*) also tended to be similar across a range of data filtering schemes (Fig. S4–6). Instead, we suggest a different risk associated with amplification biases: because the signal for gene flow is expected to be stronger in smaller chromosomes(Martin et al. 2019), biased sequencing experiments, including ours, which preferentially sequence regions with high GC content on smaller chromosomes could artificially inflate genome-wide estimates for gene flow. We thus suggest estimates of introgression inferred from RADseq datasets be interpreted with caution.

**SUPPLEMENTAL TABLES:**

| **Institution** | **Voucher** | **Tissue_No** | **CollectNo** | **Sex** | **Sample** | **Latitude** | **Longitude** | **Population** |
| --- | --- | --- | --- | --- | --- | --- | --- | --- |
| AMNH | NA | J249 | GDR233 | Male | T_ae_J249_ma | -7.56722 | -60.68817 | S. Rondonia |
| AMNH | NA | J252 | GDR234 | Female | T_ae_J252_ma | -7.56722 | -60.68817 | S. Rondonia |
| AMNH | NA | J261 | ROS006 | Unspecified | T_ae_J261_ma | -7.56722 | -60.68817 | S. Rondonia |
| AMNH | NA | J298 | GDR247 | Male | T_ae_J298_ma | -7.56722 | -60.68817 | S. Rondonia |
| AMNH | NA | J319 | LJM225 | Male | T_ae_J319_roar | -7.699787 | -60.890504 | S. Rondonia |
| AMNH | NA | J419 | GDR283 | Female | T_ae_J419_roar | -7.699787 | -60.890504 | S. Rondonia |
| AMNH | NA | J598 | MAR131 | Unspecified | T_ae_J598_arsu | -7.703667 | -60.595694 | N. Rondonia |
| AMNH | NA | J616 | GT141 | Female | T_ae_J616_arsu | -7.703667 | -60.595694 | N. Rondonia |
| AMNH | NA | J678 | GDR370 | Female | T_ae_J678_roar | -8.17761 | -60.45314 | S. Rondonia |
| INPA | NA | A15069 | CD 238 | Female | T_ae_A15069_suta | -3.316667 | -55.333333 | N. Rondonia |
| INPA | NA | A16041 | CD 356 | Female | T_ae_A16041_suta | -4.5 | -56.3 | N. Rondonia |
| INPA | NA | A2720 | EMP 15 | Male | T_ae_A2720_pu | -3.683333 | -60.316667 | E. Inambari |
| INPA | NA | A2833 | EB 44 | Male | T_ae_A2833_pu | -4.983333 | -61.566667 | E. Inambari |
| INPA | NA | A317 | AMF 153 | Male | T_ae_A317_jigu | -8.853298 | -63.86298 | S. Rondonia |
| INPA | NA | A324 | AMFP 5 | Female | T_ae_A324_jigu | -8.853298 | -63.86298 | S. Rondonia |
| INPA | NA | A3559 | EB 135 | Male | T_ae_A3559_pu | -8.840933 | -64.061614 | E. Inambari |
| INPA | NA | A439 | AMFP 46 | Female | T_ae_A439_pu | -5.241667 | -60.716667 | E. Inambari |
| INPA | NA | A458 | CLB 36 | Male | T_ae_A458_ma | -6.3 | -60.4 | S. Rondonia |
| INPA | NA | A509 | AMFP 72 | Male | T_ae_A509_ma | -6.3 | -60.4 | S. Rondonia |
| INPA | NA | A522 | ITM 17 | Male | T_ae_A522_ma | -6.3 | -60.333333 | N. Rondonia |
| INPA | NA | A7968 | GRL 927 | Male | T_ae_A7968_In | -4.983333 | -62.966667 | W. Inambari |
| INPA | NA | A8087 | GRL 961 | Female | T_ae_A8087_In | -4.983333 | -62.966667 | W. Inambari |
| LSUMNS | NA | B-28095 | NA | Unspecified | T_ae_28095_In | -7.133333 | -75.683333 | W. Inambari |
| LSUMNS | NA | B-75520 | NA | Unspecified | T_ae_75520_In | -10.38 | -73.72 | W. Inambari |
| LSUMNS | NA | B-80508 | NA | Unspecified | T_ae_80508_arsu | -5.288889 | -59.693611 | N. Rondonia |
| LSUMNS | NA | B-80716 | NA | Unspecified | T_ae_80716_arsu | -5.25175 | -59.694722 | N. Rondonia |
| LSUMNS | NA | B-81278 | NA | Female | T_ae_81278_arsu | -4.050833 | -59.115278 | N. Rondonia |
| LSUMNS | NA | B-81338 | NA | Unspecified | T_ae_81338_arsu | -4.050833 | -59.115278 | N. Rondonia |
| LSUMNS | NA | B-85274 | NA | Unspecified | T_ae_85274_suta | -5.797861 | -59.230778 | N. Rondonia |
| LSUMNS | NA | B-86147 | NA | Female | T_ae_86147_arsu | -6.754722 | -59.083056 | N. Rondonia |
| LSUMNS | NA | B-86229 | NA | Female | T_ae_86229_arsu | -6.751667 | -59.075556 | N. Rondonia |
| LSUMNS | NA | B-86569 | NA | Unspecified | T_ae_86569_arsu | -6.562222 | -59.093889 | N. Rondonia |
| MPEG | NA | T82491 | PUR008 | Unspecified | T_ae_82491_pu | -7.69452 | -65.223491 | E. Inambari |
| MPEG | NA | T82606 | PUR124 | Unspecified | T_ae_82606_pu | -7.336479 | -65.15999 | W. Inambari |
| MPEG | NA | T82608 | PUR126 | Female | T_ae_82608_pu | -7.336479 | -65.15999 | W. Inambari |
| MPEG | 67048 | T10229 | FPR085 | Male | T_ae_T10229_suta | -3.946111 | -58.456111 | N. Rondonia |
| MPEG | 68883 | T12313 | TUP057 | Male | T_ae_T12313_pu | -4.083333 | -60.6605 | E. Inambari |
| MPEG | 71057 | T13219 | OM168 | Female | T_ae_T13219_pu | -7.518333 | -63.337778 | E. Inambari |
| MPEG | 71058 | T13278 | OM228 | Male | T_ae_T13278_ma | -8.909139 | -62 | S. Rondonia |
| MPEG | 72839 | T14205 | AMA341 | Male | T_ae_T14205_In | -4.530278 | -71.61625 | W. Inambari |
| MPEG | 72357 | T14601 | ARA075 | Female | T_ae_T14601_suta | -2.783333 | -55.6 | N. Rondonia |
| MPEG | 74122 | T16625 | ARAII029 | Male | T_ae_T16625_suta | -3.094167 | -55.535639 | N. Rondonia |
| MPEG | 74194 | T16704 | ARAII102 | Male | T_ae_T16704_suta | -2.586833 | -55.195167 | N. Rondonia |
| MPEG | 76369 | T19436 | JAT(C)269 | Male | T_ae_T19436_suta | -5.073075 | -56.8559 | N. Rondonia |
| MPEG | 58705 | T2166 | MPDS665 | Female | T_ae_T2166_ma | -7.466667 | -62.816667 | S. Rondonia |
| MPEG | 58706 | T2207 | MPDS720 | Male | T_ae_T2207_ma | -7.55 | -62.55 | S. Rondonia |
| MPEG | 54953 | T3237 | OP115 | Male | T_ae_T3237_jigu | -10.833333 | -64.75 | S. Rondonia |
| MPEG | 54955 | T3260 | OP150 | Female | T_ae_T3260_jigu | -10.833333 | -64.75 | S. Rondonia |
| MPEG | 60171 | T3376 | CUJ133 | Male | T_ae_T3376_jigu | -10.833333 | -64.75 | W. Inambari |
| MPEG | 58707 | T4355 | MPDS721 | Female | T_ae_T4355_ma | -7.55 | -62.55 | S. Rondonia |
| MPEG | 58100 | T742 | JRT060 | Female | T_ae_T742_suta | -2.6 | -56.183333 | N. Rondonia |

**Table S1:** A list of samples used in this study.

| **Sliding Window Size** | **Minimum number of SNPs** | **Number of windows** |
| --- | --- | --- |
| 5000 | 5 | 2054 |
| 5000 | 10 | 124 |
| 10000 | 5 | 3063 |
| 10000 | 10 | 336 |
| **50000** | **5** | **4858** |
| 50000 | 10 | 1727 |
| 100000 | 5 | 4567 |
| 100000 | 10 | 2181 |
| 200000 | 5 | 3525 |
| 200000 | 10 | 2205 |

**Table S2:** The number of windows in our dataset fighting criteria for window size and minimum number of SNPs. We chose 50,000 bp sliding windows with a minimum of 5 SNPs to use for generating gene trees across the genome.

| **chromosome** | **No. gene trees** | **chromosome** | **No. gene trees** |
| --- | --- | --- | --- |
| **1** | 328 | **18** | 163 |
| **2** | 322 | **19** | 129 |
| **3** | 312 | **20** | 118 |
| **4** | 253 | **21** | 154 |
| **5** | 230 | **22** | 87 |
| **6** | 245 | **23** | 105 |
| **7** | 152 | **24** | 94 |
| **8** | 193 | **25** | 100 |
| **9** | 210 | **26** | 72 |
| **10** | 167 | **27** | 79 |
| **11** | 164 | **28** | 50 |
| **12** | 159 | **29** | 29 |
| **13** | 153 | **30** | 23 |
| **14** | 155 | **31** | 5 |
| **15** | 172 | **32** | 1 |
| **16** | 161 | **W** | 5 |
| **17** | 175 | **Z** | 83 |

**Table S3:** The number of genetrees (based on 50kb windows with a minimum of 5 SNPs) that were reconstructed for each chromosome.

| **Parameters** | **Estimates** | **SD** | **R^2^** | **MAE** |
| --- | --- | --- | --- | --- |
| Ne W Inambari (WI) | 1,106,527 | 53,933 | 0.939 | 70,793 |
| Ne E Inambari (EI) | 198,314 | 58,852 | 0.93 | 72,272 |
| Ne N Rondonia (NR) | 1,120,729 | 42,098 | 0.94 | 68,653 |
| Ne S Rondonia (SR) | 843,112 | 129,902 | 0.948 | 65,283 |
| Tdiv (WI,SR) | 149,546 | 6,010 | 0.948 | 48,492 |
| Tdiv (SR,(WI,EI)) | 616,206 | 54,350 | 0.614 | 117,941 |
| Tdiv (NR,(SR,WI,EI)) | 824,957 | 106,470 | 0.61 | 125,875 |
| mig SR<->EI | 3.496 | 0.187 | 0.935 | 0.224 |
| mig SR<->NR | 3.349 | 0.096 | 0.956 | 0.181 |

**Table S4:** Demographic parameter estimates obtained with a Neural Network regression approach. Ne - population effective size in diploid individuals; Tdiv - Divergence times in years ago; mig - migratio in individuals per generation; Absolute estimates and standard deviation (SD) are based on the average for 10 replicates; R^2^ values are based on a linear regression between simulated and estimated values, and represent the overall confidence on the parameter estimation; MAE - Mean absolute error.

**Supplemental Figures:**

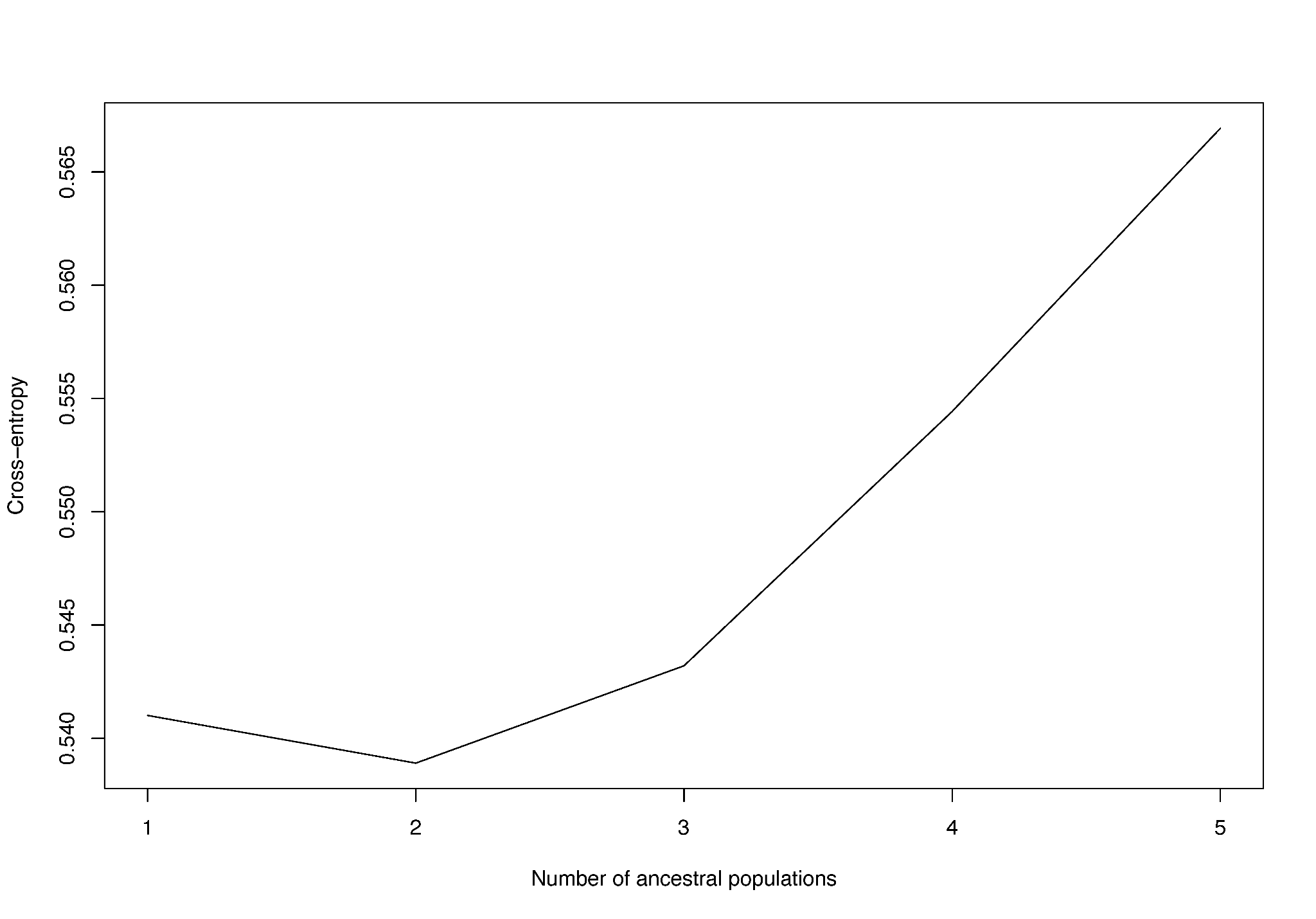

**Figure S1:** Cross entropy plot showing results from sNMF runs for K=1–5. K=2 has the lowest cross entropy and is the best-fit K-value.

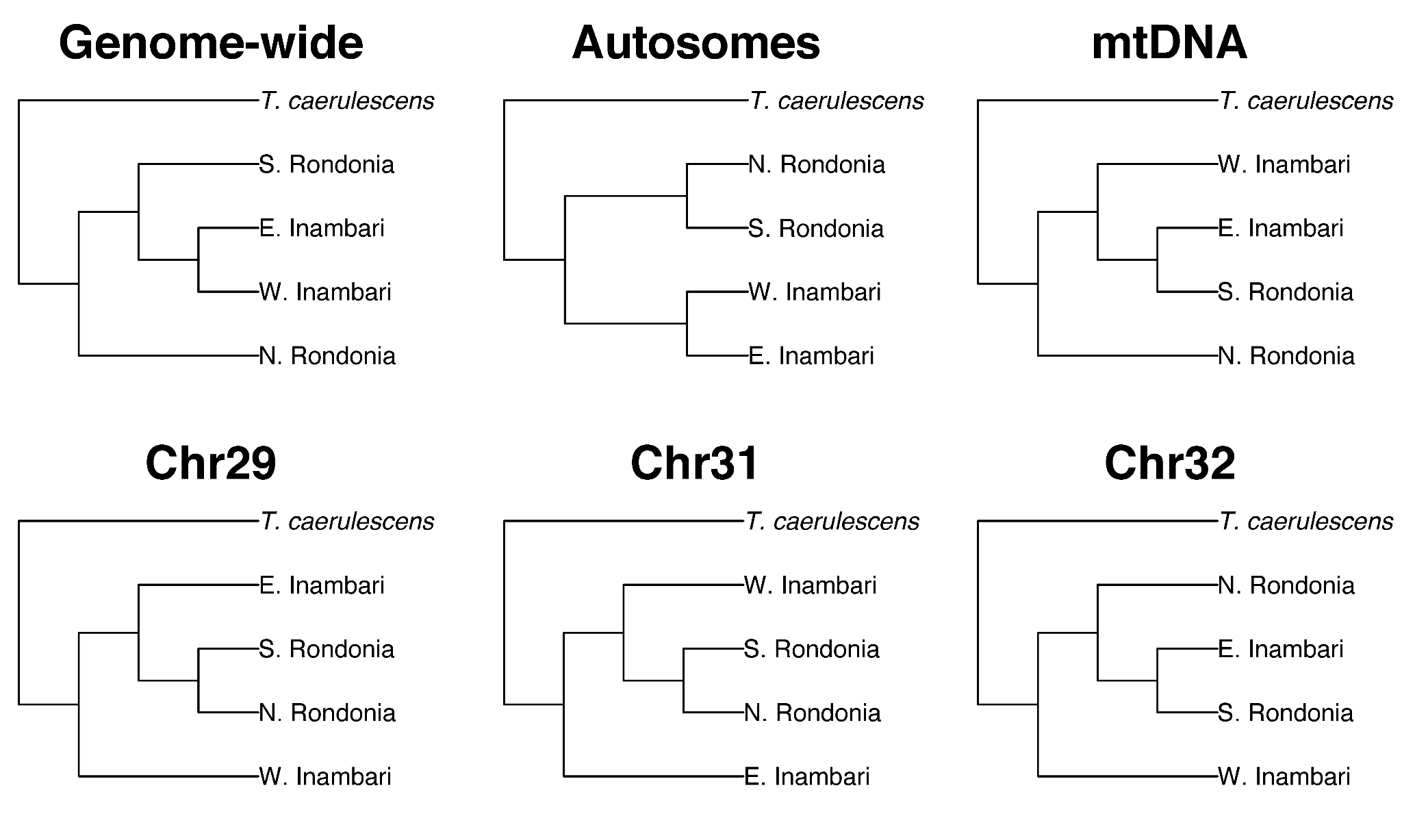

**Figure S2:** ASTRAL results for various portions of the genome including all loci combined (genome-wide), all autosomal loci combined (autosomes), and the topologies for three chromosomes not shown in the main text (Chr29, Chr31, and Chr32). The mitochondrial topology from a previous study is also shown (mtDNA).

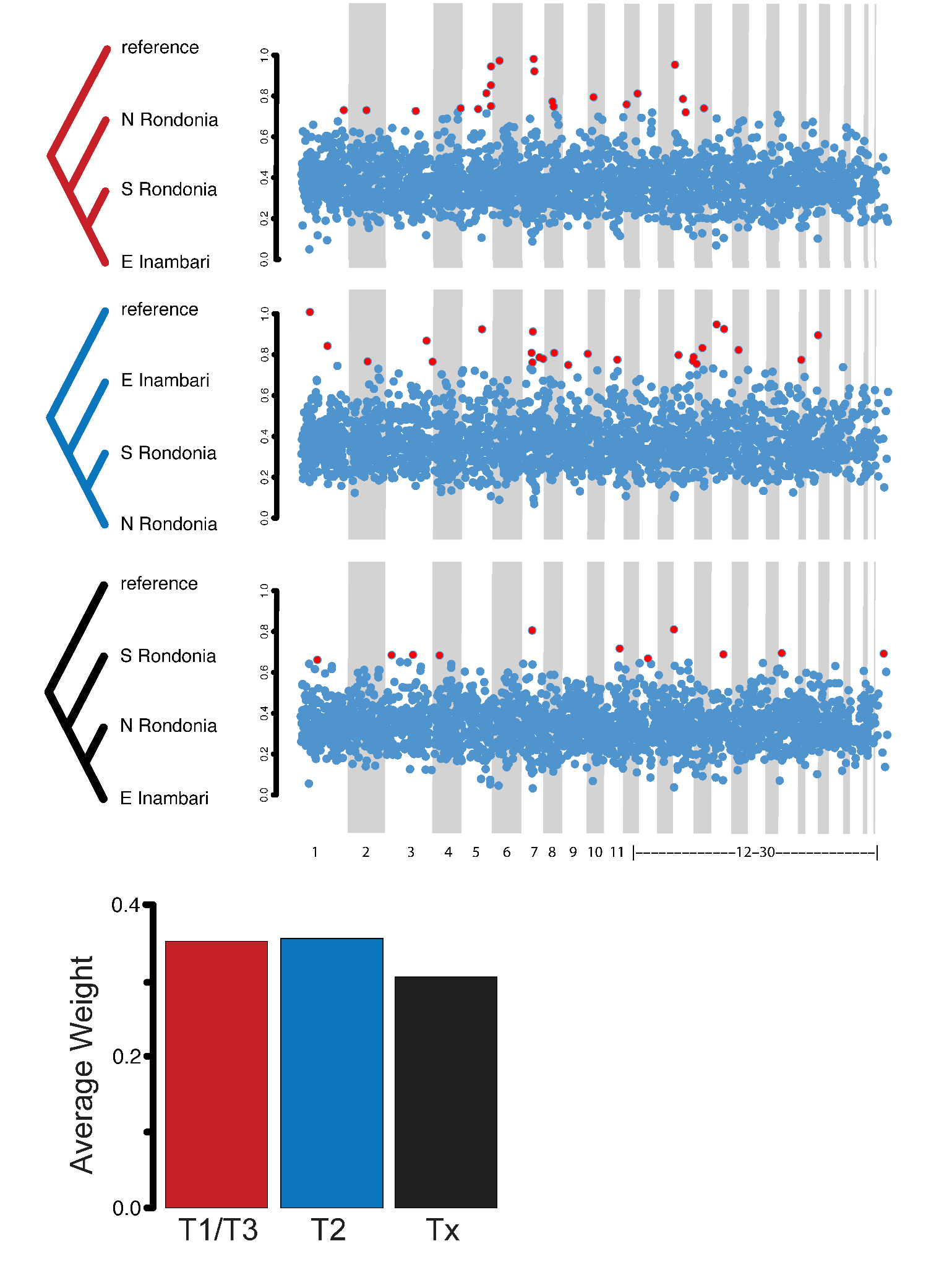

**Figure S3:** Weights for alternative unrooted topologies (W Inambari population dropped) across the autosomes, including T1/T3 (A), T2 (B), and a third unrooted topology, Tx (C). Red points represent outlier weights indicating high support for a given topology. Rooted versions of each topology are shown at the left. Average autosome-wide weights for each topology are shown for TWISST analysis including (D) and excluding the reference (E) genome as an outgroup. Because these topologies are unrooted, T1 is indistinguishable from T3 in analyses that include the reference, and T1 is indistinguishable from T2 when the reference is omitted.

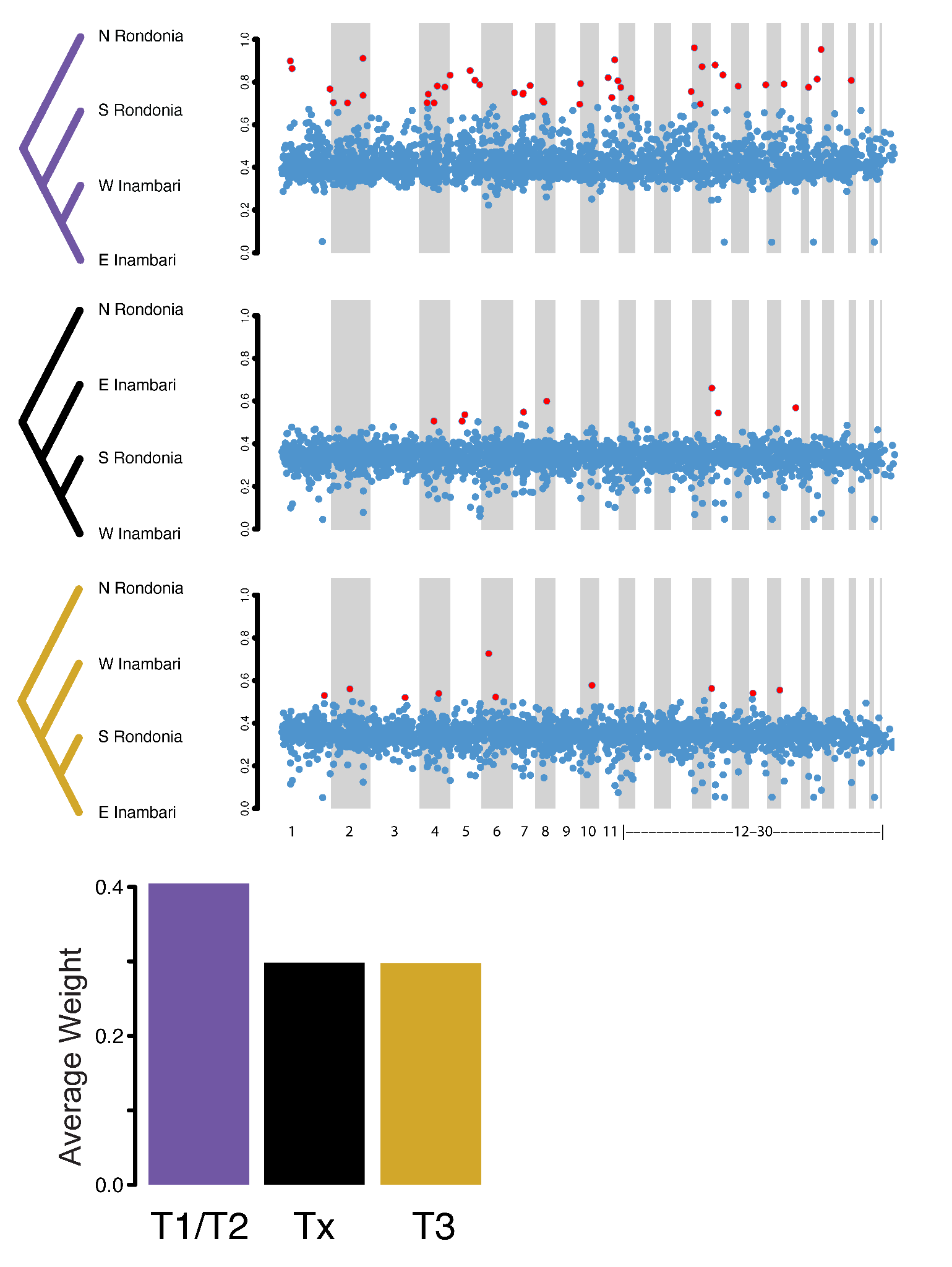

**Figure S4:** Weights for alternative unrooted topologies for ingroup populations across the autosomes. (A) Rooted versions of the topology and their weights for windows across the genome (B). Red points represent outlier weights indicating high support for a given topology.

(C) The genome-wide average is shown at the right.

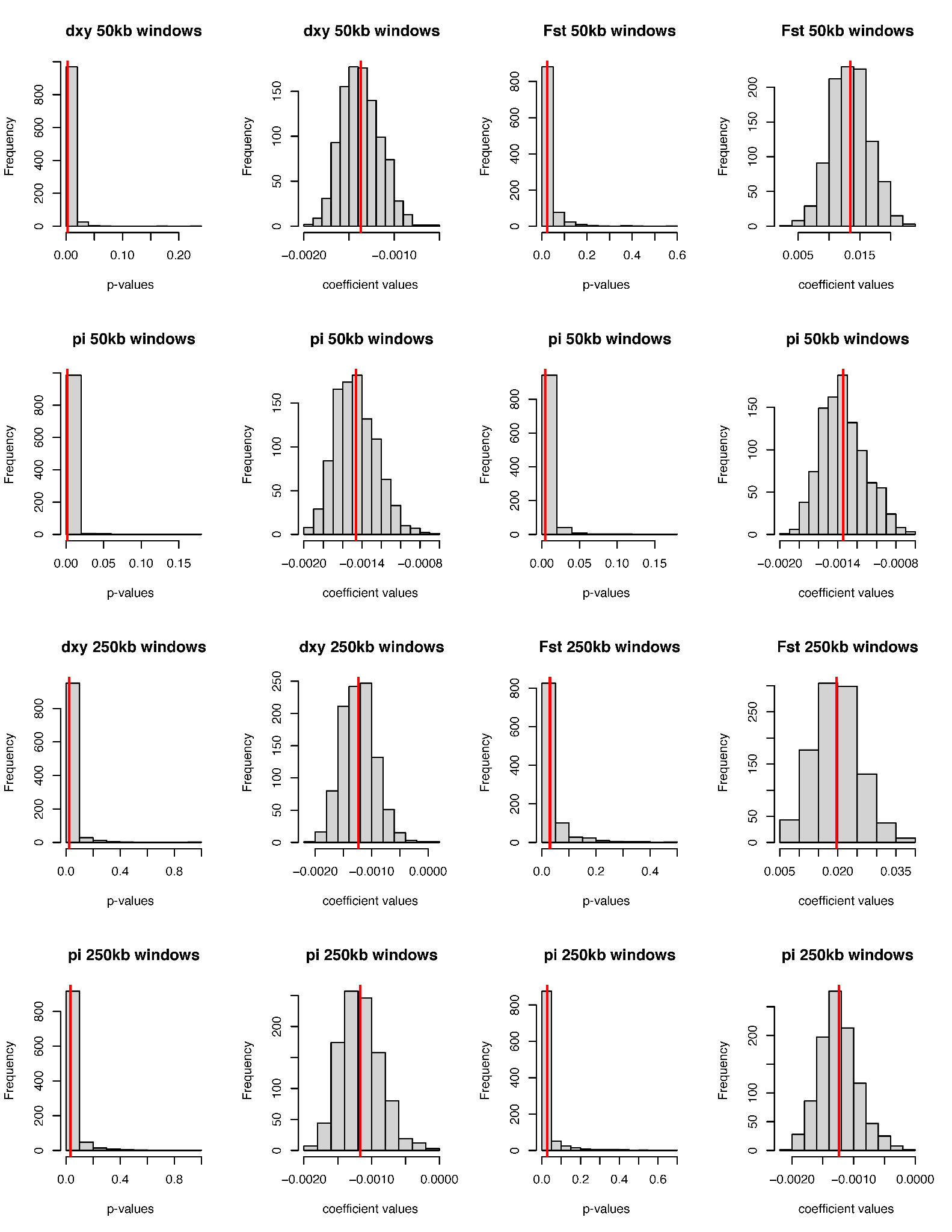

**Figure S5:** Results of the bootstrapping exercise on models explaining d_xy_, F_ST_, and π as functions of chromosome length (Figure 4). Histograms represent the distribution of p-values and linear regression coefficients across bootstrap replicates. Vertical red lines indicate the mean value. Mean p-values are all significant (*P*<0.05).

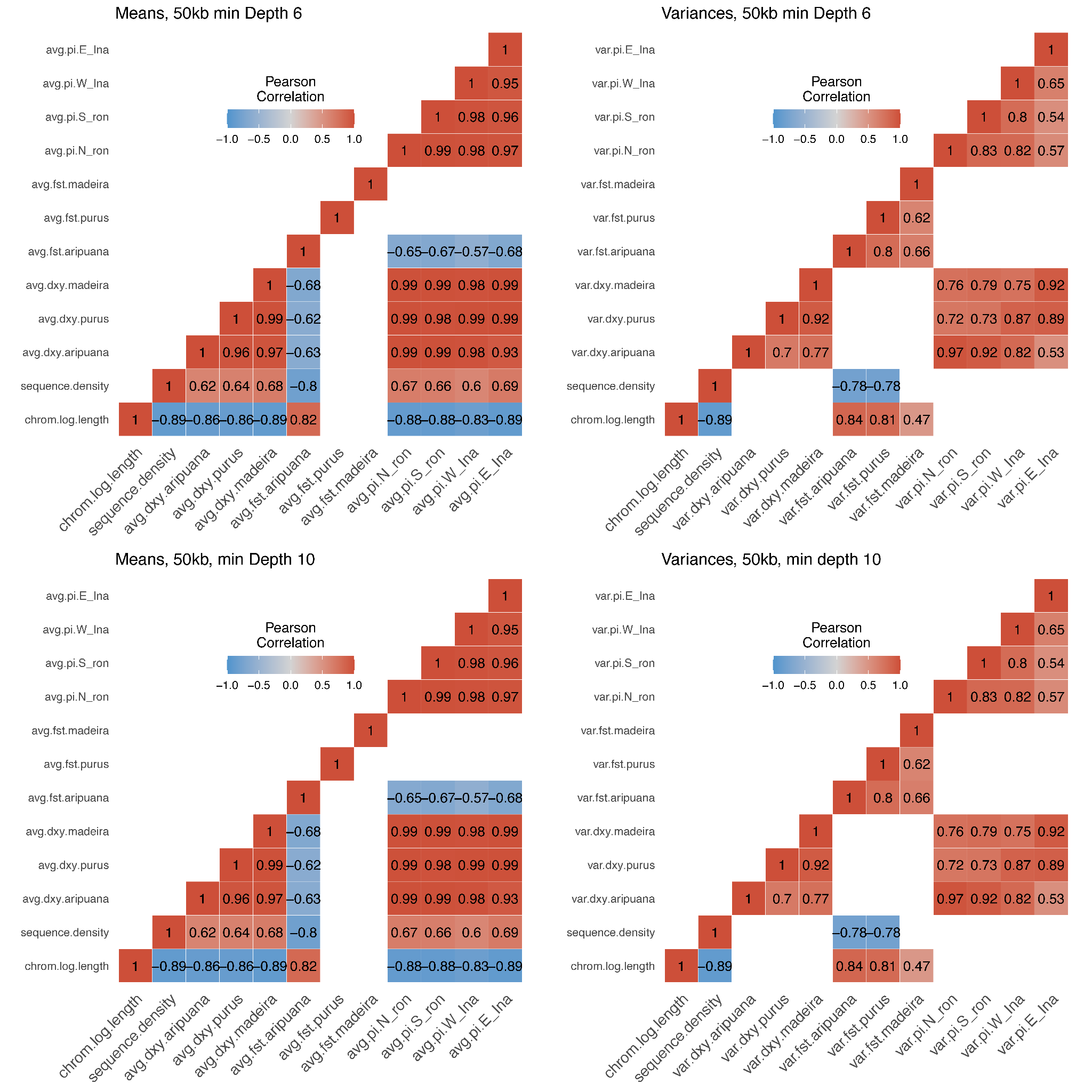

**Figure S6:** Heatmap of Pearson’s coefficients for pairwise correlations of genetic summary statistics for chromosomes for different VCF file filtering schemes. The heatmaps at the left show pairwise correlations of within-chromosome means, and the heatmaps at the right show within-chromosome variances for values of d_xy_, F_ST_, and pi . Empty (white) boxes represent correlations with nonsignificant P-values (*P*>0.05).

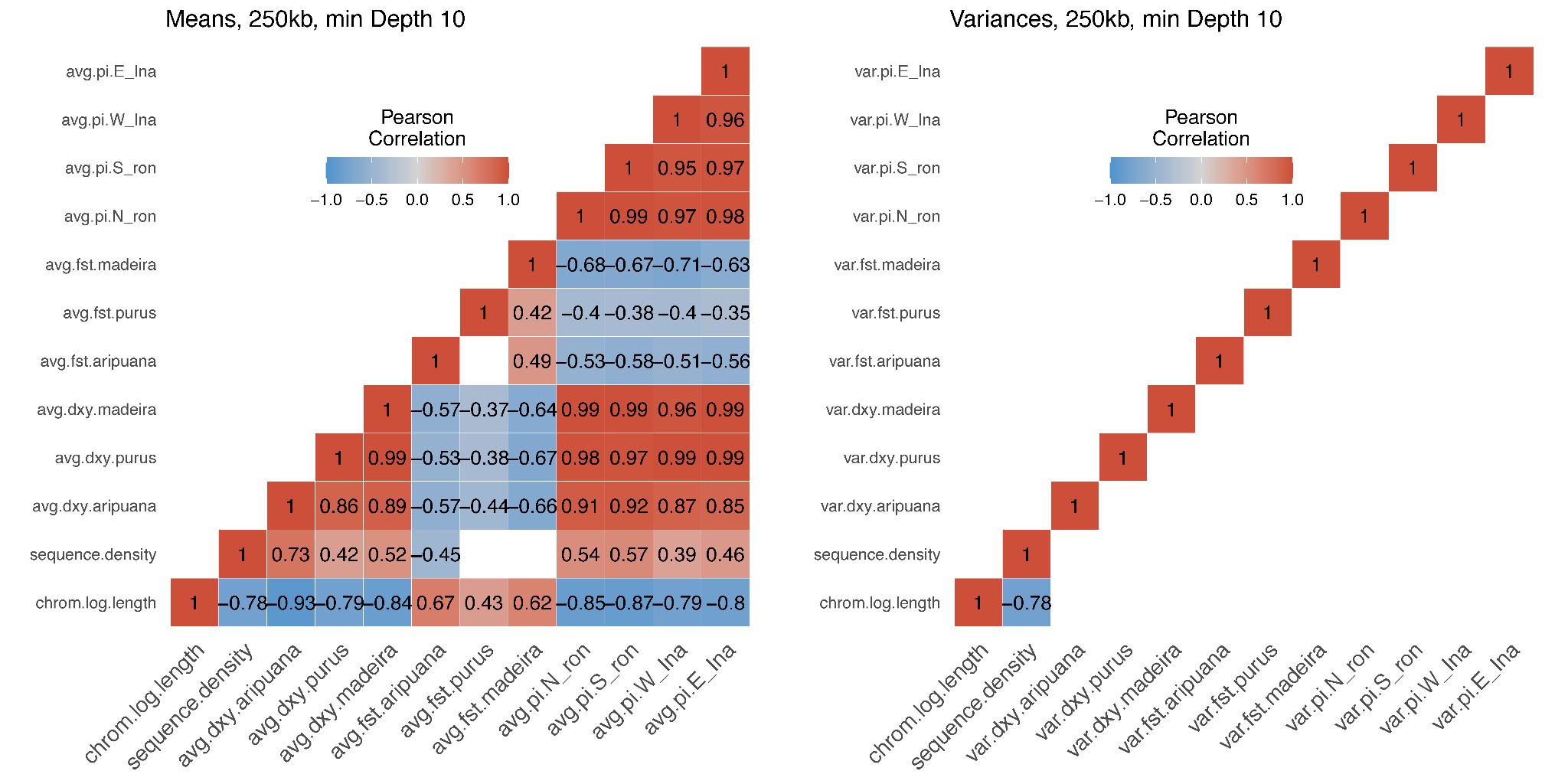

**Figure S7:** Heatmap of Pearson’s coefficients for pairwise correlations of genetic summary statistics for chromosomes for different VCF file filtering schemes. The heatmaps at the left show pairwise correlations of within-chromosome means, and the heatmaps at the right show within-chromosome variances for values of d_xy_, F_ST_, and pi . Empty (white) boxes represent correlations with nonsignificant P-values (*P*>0.05).

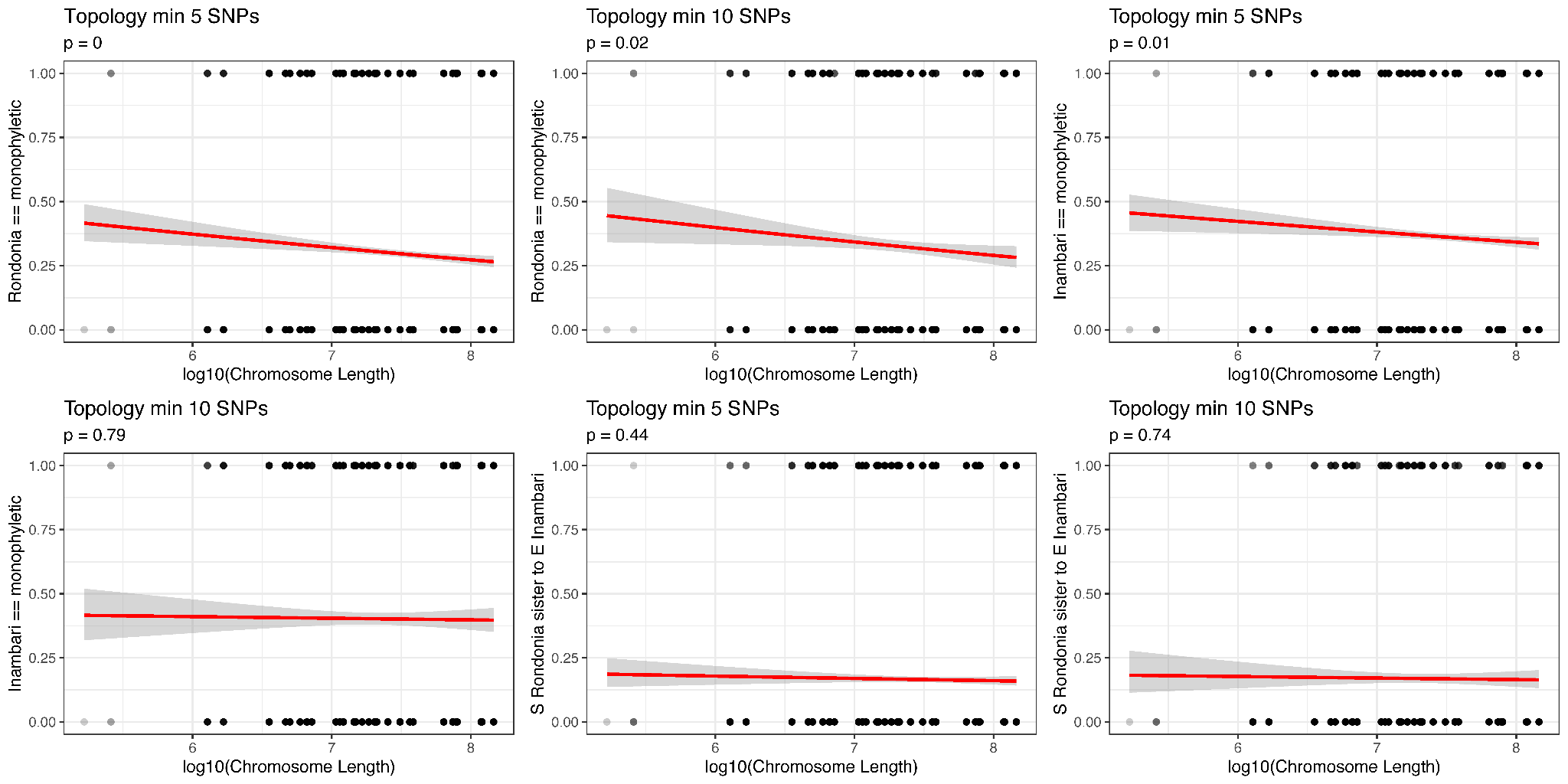

**Figure S8:** Logistic regressions for genetree topology and chromosome length. We show results for gene trees with a minimum of 5 SNPs and genetrees with a minimum of 10 SNPs.

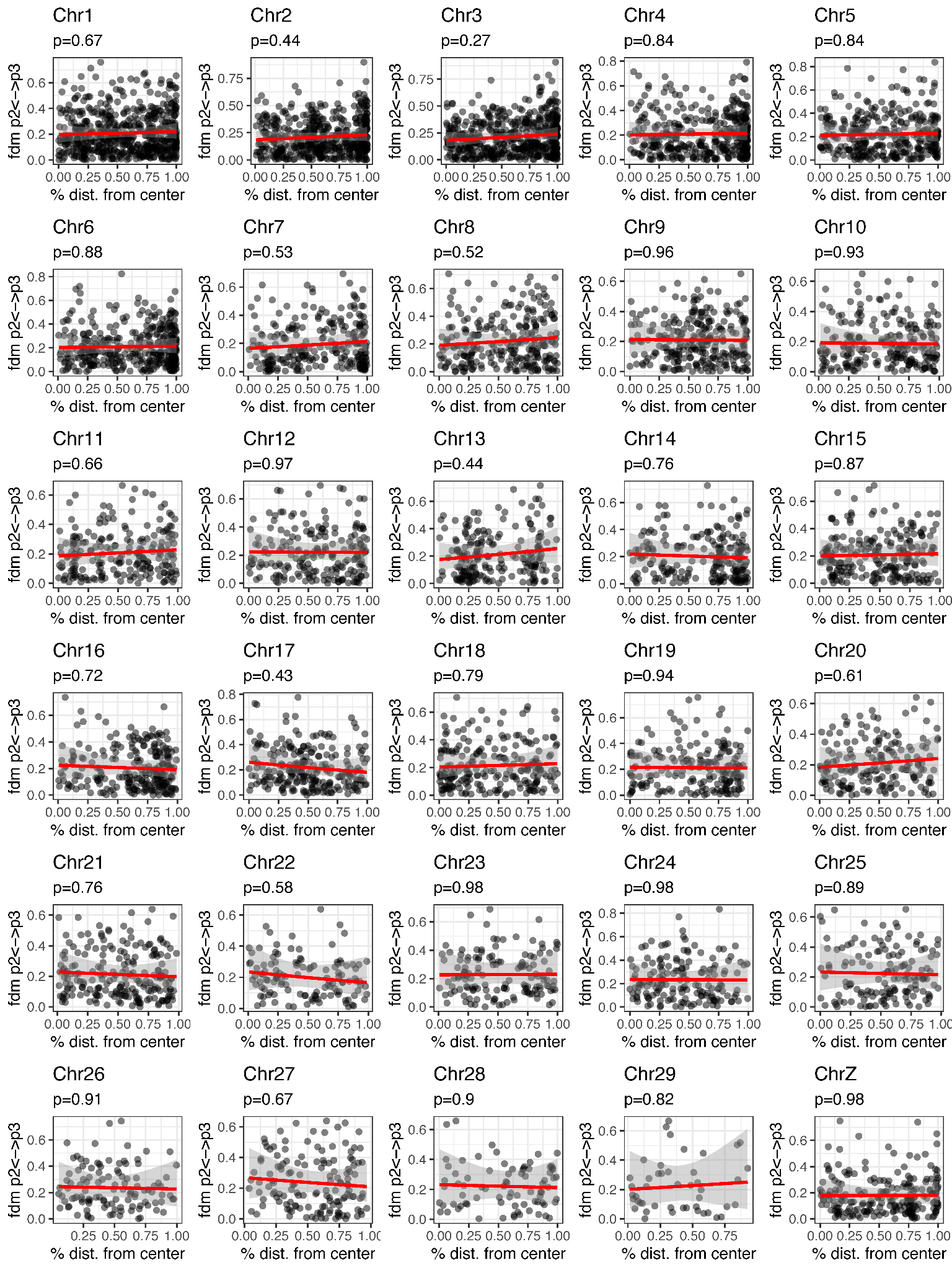

**Figure S9**: Linear regressions testing for an association between introgression and distance from chromosome center.

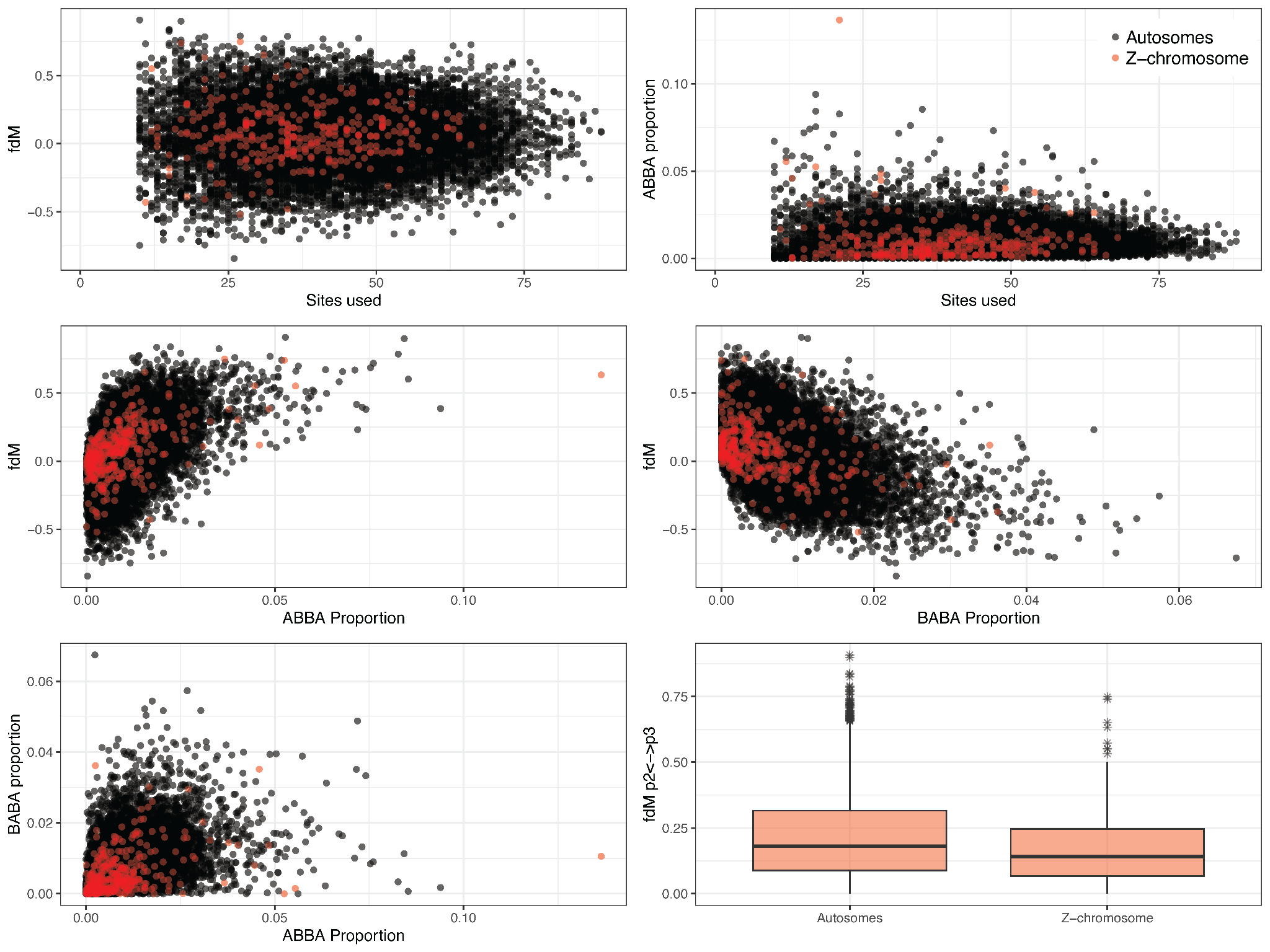

**Figure S10:** Scatterplots showing variation in introgression statistics across the genome. In the top row, variance in values for *𝑓*_dM_ is relatively uniformly distributed across the genome and not affected by sliding window content (sites used). Overall however, windows on the Z-chromosome had lower rates of introgression than those from the autosomes (bottom right).

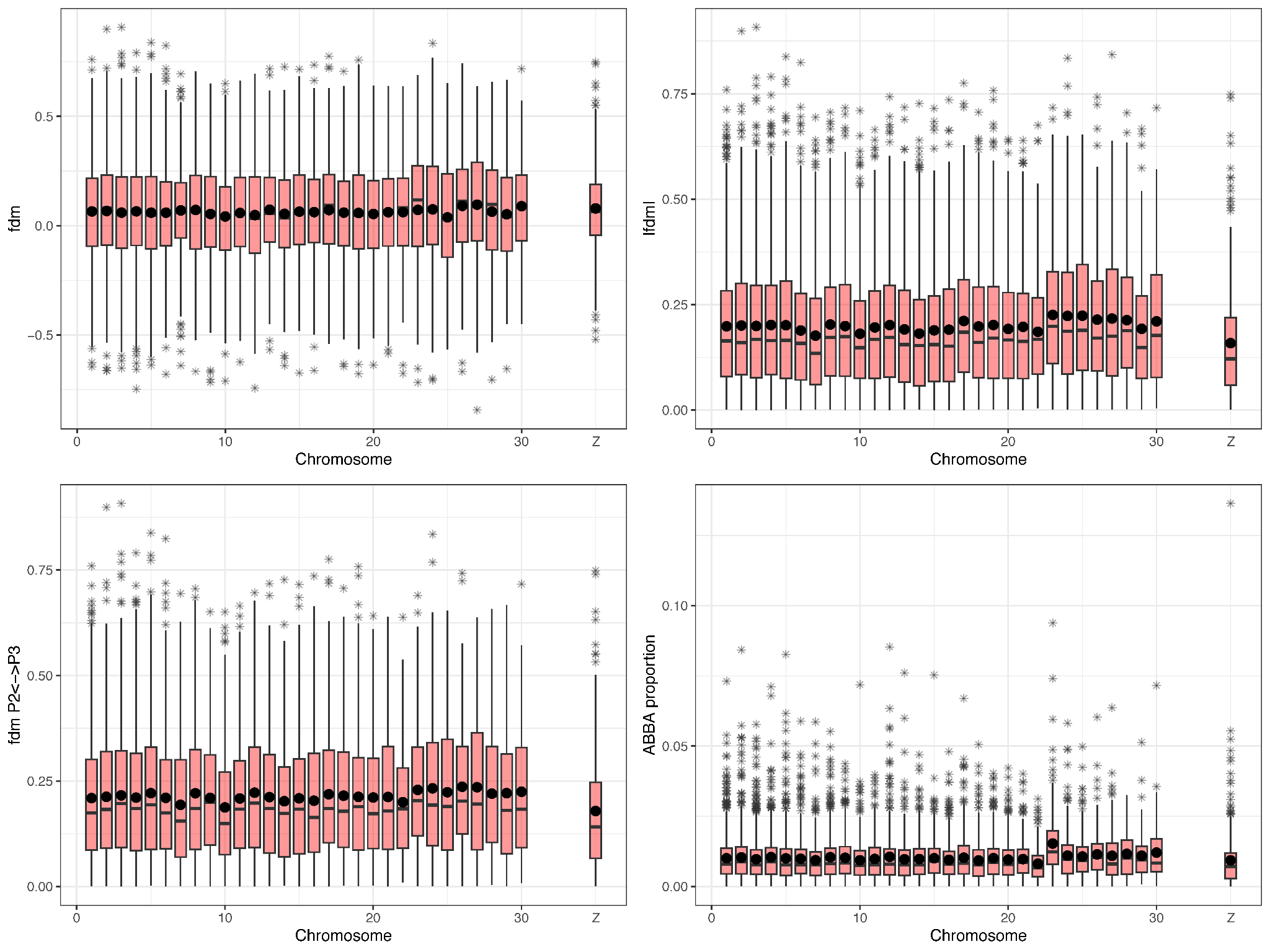

**Figure S11:** Estimated introgression statistics across the genome. Top left: boxplot showing *𝑓*_dM_ values for each chromosome. Top right: boxplot of the absolute value of *𝑓*_dM_ for each chromosome. Bottom left: boxplot showing the positive values of *𝑓*_dM_ only per chromosome. Bottom right: boxplots showing the proportion of ABBA’s per window for each chromosome. Boxplots show the median (black bars), mean (black points).

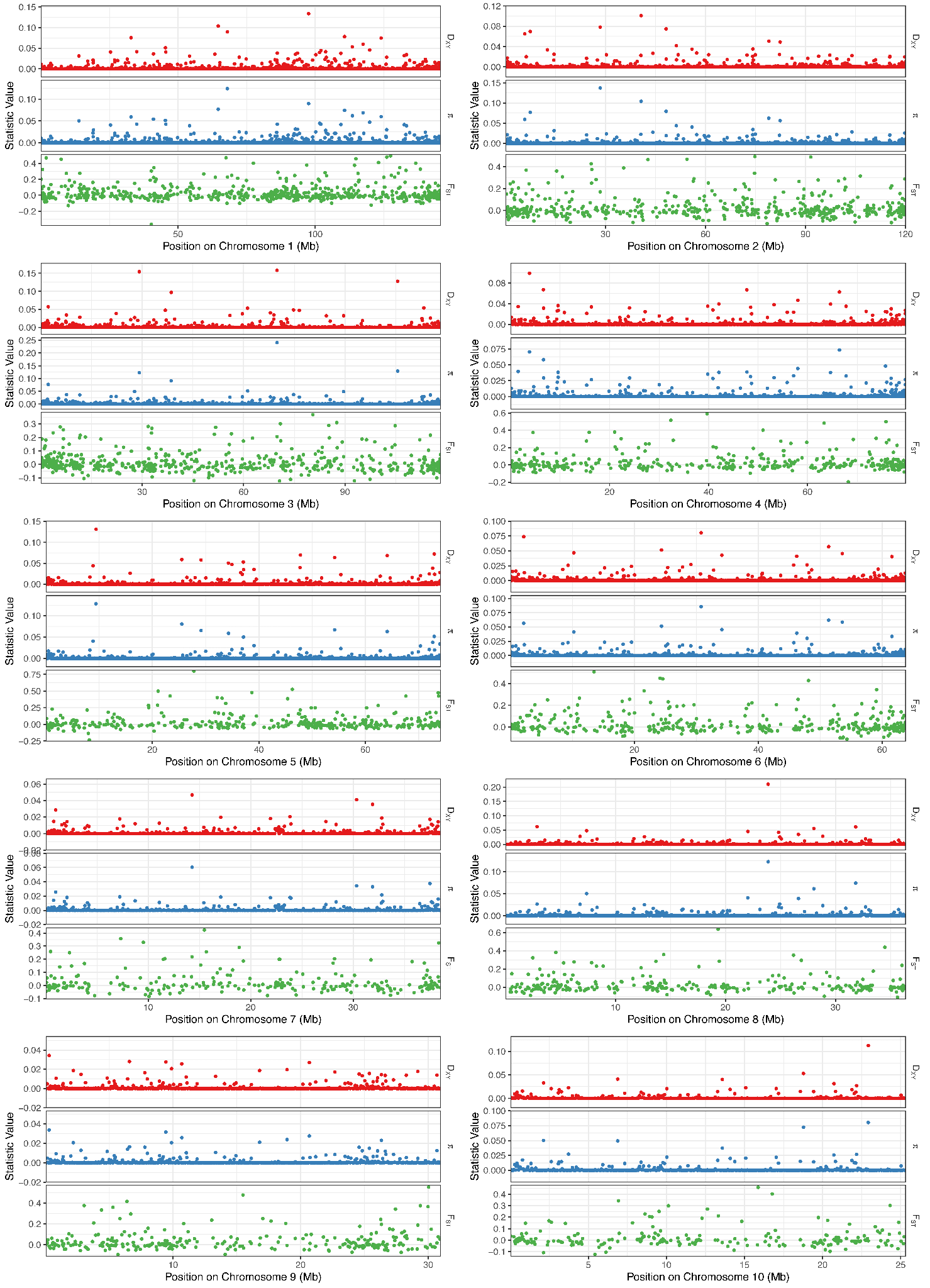

**Figure S12:** Plotted summary statistics for 50 kb sliding windows across the genome (chromosomes 1–10).

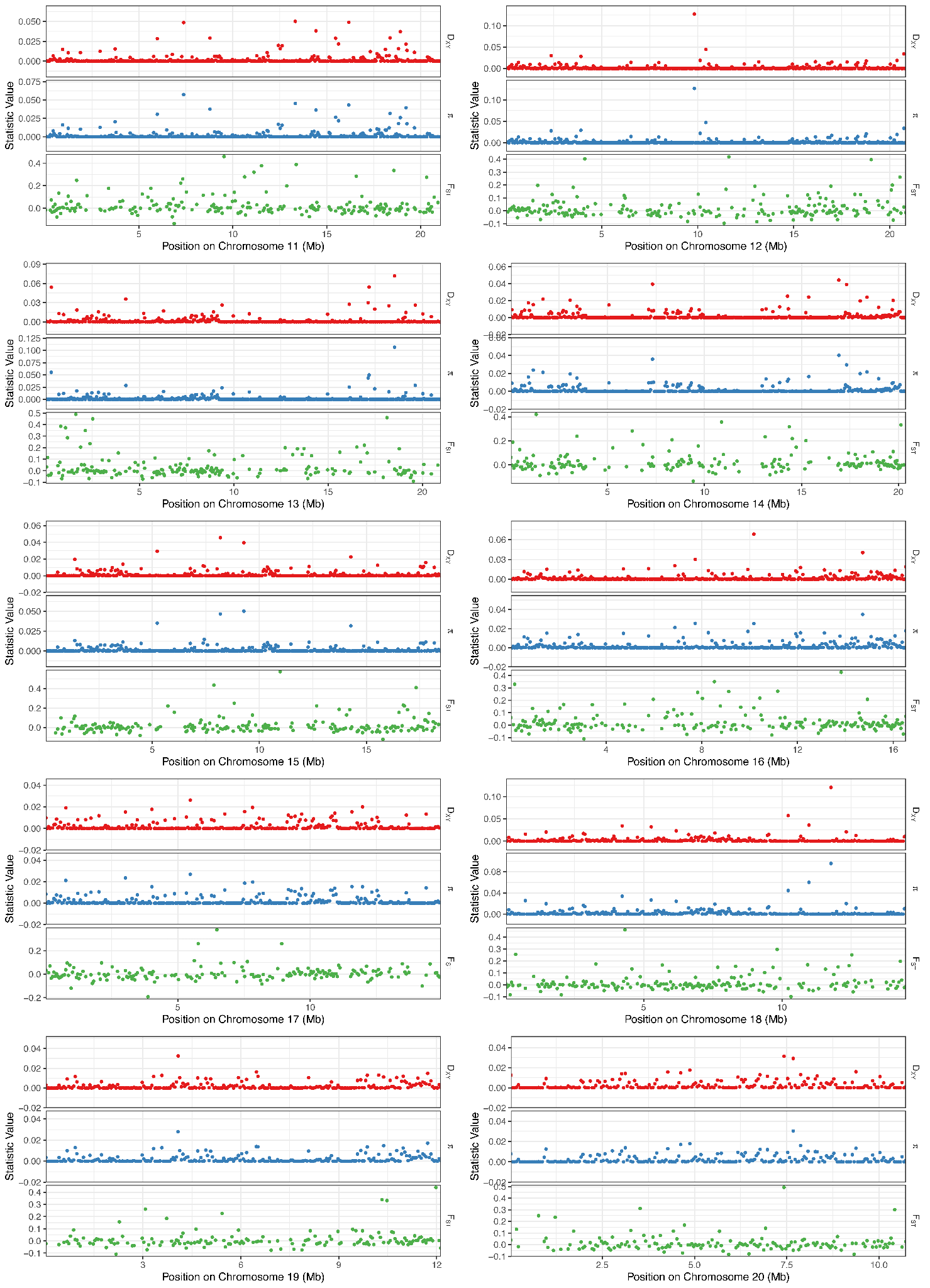

**Figure S13:** Plotted summary statistics for 50 kb sliding windows across the genome (chromosomes 11–20).

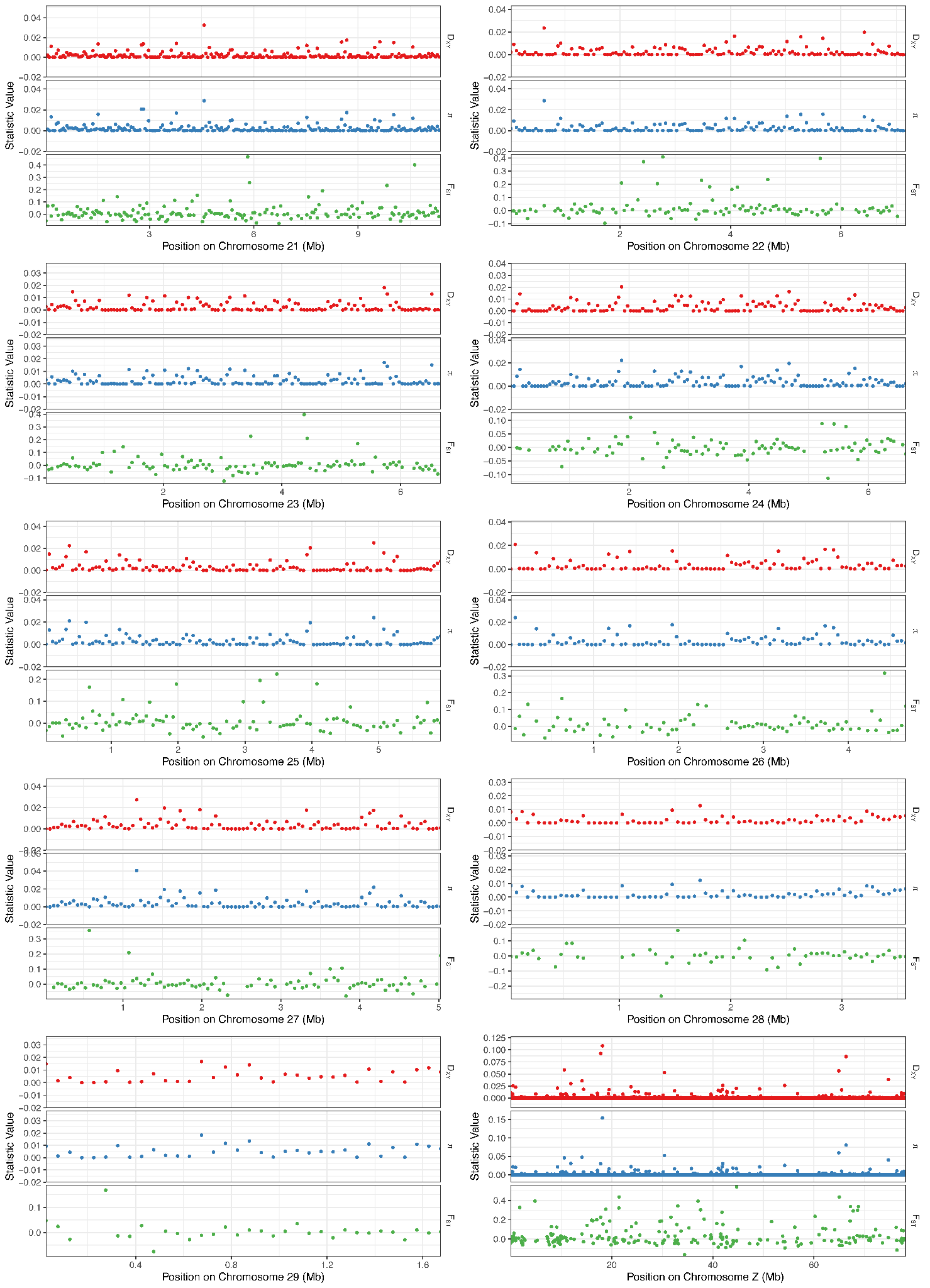

**Figure S14:** Plotted summary statistics for 50 kb sliding windows across the genome (chromosomes 21–29 and Z).
